# Pan-cancer genomic analysis shows hemizygous *PTEN* loss tumors are associated with immune evasion and poor outcome

**DOI:** 10.1101/2022.09.16.508308

**Authors:** T Vidotto, CM Melo, W Lautert-Dutra, LP Chaves, RB Reis, JA Squire

**Author notes:** Corresponding author: Jeremy Andrew Squire. Joint first authors (T Vidotto, CM Melo).

## Abstract

In tumors, somatic mutations of the *PTEN* suppressor gene are associated with advanced disease, chemotherapy resistance, and poor survival. *PTEN* loss of function may occur by inactivating mutation, by deletion, either affecting one copy (hemizygous loss) leading to reduced gene expression or loss of both copies (homozygous) with expression absent. Various murine models have shown that minor reductions in PTEN protein levels strongly influence tumorigenesis. Most *PTEN* biomarker assays dichotomize PTEN (i.e. presence vs. absence) ignoring the role of one copy loss. We performed a *PTEN* copy number analysis of 9,793 TCGA cases from 30 different tumor types. There were 419 (4.28%) homozygous and 2484 (25.37%) hemizygous *PTEN* losses. Hemizygous deletions led to reduced PTEN gene expression, accompanied by increased levels of instability and aneuploidy across tumor genomes. Outcome analysis of the pan-cancer cohort showed that losing one copy of *PTEN* reduced survival to comparable levels as complete loss, and was associated with transcriptomic changes controlling immune response and the tumor microenvironment. Immune cell abundances were significantly altered for *PTEN* loss, with changes in head and neck, cervix, stomach, prostate, brain, and colon more evident in hemizygous loss tumors. These data suggest that reduced expression of *PTEN* in tumors with hemizygous loss leads to tumor progression and influences anticancer immune response pathways.

## INTRODUCTION

The PTEN tumor suppressor functions as a phosphatase, which metabolizes PIP3, the lipid product of PI3-Kinase that antagonizes the activation of the oncogenic PI3K/AKT/mTOR signaling axis ^1,2^. In addition to its role as a primary regulator of cancer cell growth, recent findings also implicate PTEN in mediating aspects of the immune response against cancer ^3,4^.

*PTEN* is frequently lost, either partially affecting one copy of the gene (hemizygous loss; HemDel) or entirely affecting both copies of the gene (homozygous loss; HomDel), in a range of sporadic tumor types. A variety of mechanisms can cause *PTEN* gene inactivation, including missense and truncation mutations and deletions, as well as reduced expression of active PTEN protein mediated by promoter methylation, the effects of miRNAs, and the suppression of PTEN enzyme activity ^5^. Loss of function of the *PTEN* tumor suppressor has been observed as one of the most common events in many types of cancer ^6^, having the greatest selection pressure for deletion based on an analysis of 746 human cancer genomes ^7^.

The *Pten* conditional knockout mouse model closely mimics human prostate initiation and progression ^8^. Studies showed that loss of one copy of the *Pten* allele dramatically increases the rate of tumors ^9^. It was proposed that haploinsufficiency of the *Pten* gene resulted in reduced levels of the functional protein that were insufficient for preventing tumor formation. Similar studies using a brain-tumor-specific model also showed that *Pten* haploinsufficiency accelerated the formation of glioblastomas ^10^. Analysis of hypomorphic *Pten* mice demonstrated that as little as a 20% reduction of PTEN protein contributed to the development of prostate cancer in mice ^11^. These and other mouse studies ^12^ show that minor reductions in PTEN protein levels can profoundly influence tumorigenesis ^13^. The findings from *Pten* murine tumor models are consistent with the precise cellular controls needed to regulate *PTEN* stability and function ^14^, implying that even subtle changes in *PTEN* expression may have profound effects on cancer development and progression ^5^.

Álvarez-Garcia et al. ^15^ recently surveyed the solid tumor literature on *PTEN* loss resulting from a mutation within the coding region of the gene; hemizygous or homozygous deletions of the gene as assessed by loss of heterozygosity (LOH), reduced copy number in array CGH analysis or loss of *PTEN* by FISH. Most tumor-associated missense mutations resulted in a loss or significantly reduced phosphatase activity or protein truncations. *PTEN* was found to demonstrate distinct patterns of somatic mutation leading to loss of function across different tumor types. Most glioblastomas display loss of function of *PTEN* in agreement with data from TCGA showing *PTEN* deletions in 143/170 (85%) of glioblastomas ^16^. Other human tumor types found to have frequent *PTEN* deletions and/or loss of protein are cancers of prostate (25-50%), endometrium (50%), ovary (30-50%), lung (35-55%), breast (30-40%) and colon (10-40%) ^17^. Moreover, it was found that single-copy inactivation of *PTEN* was far more common than mutation or deletion of both copies ^15^. The PI3K/AKT/mTOR consortium study further suggested that partial copy number *PTEN* loss was associated with a less favorable outcome than complete loss across all available TCGA data ^18^. These data, together with the murine *Pten* studies of tumorigenesis, formed the rationale for this PanCancer study that aimed to address the idea that partial loss of function of *PTEN* by hemizygous loss may confer a greater selective advantage for a subset of tumors than complete loss of function by homozygous loss (reviewed in ^19^).

Our previous study examining the genomic impact of *PTEN* loss using TCGA data demonstrated that primary prostate tumors harboring *PTEN* deletions had enhanced levels of aneuploidy and non-synonymous mutations ^20^. However, it was unclear whether the genomic instability generated by loss of one copy of *PTEN* was associated with tumor progression. Other studies examining the impact of *PTEN* loss using TCGA and other public domain datasets do not distinguish between one copy loss and complete loss of the gene ^21–24^. Similarly, most *PTEN* cancer biomarker studies using immunohistochemistry and fluorescence *in situ* hybridization (FISH) have not been rigorously standardized ^15,17,25,26^. For these reasons, the impact of partial loss of function due to one copy loss of *PTEN* is unknown for most cancers.

This study analyzed the TCGA copy-number and transcriptomic data, comparing oncogenic and immunophenotypic features that distinguish hemizygous *PTEN* loss from homozygous loss across the pan-cancer cohort. Our analysis also examines the relationships between *PTEN* intact, hemi-, and homozygous loss with genomic features of tumor progression and disease outcome.

## RESULTS

The *PTEN* study group comprised 30 tumor types, including 9,793 cases analyzed from the TCGA Pan-cancer cohort. *PTEN* somatic copy number alterations (SCNA) were present in 3,619 (36.96%) tumors. Copy number deletions were observed in 2,903 (29.65%) tumors, with 2,484 (25.37%) having hemizygous deletions resulting in one copy and 419 (4.28%) having homozygous loss of both copies. Copy number gains as a duplication of the *PTEN* gene were evident in 716 (7.31%) tumors (Table 1).

**Table 1.**
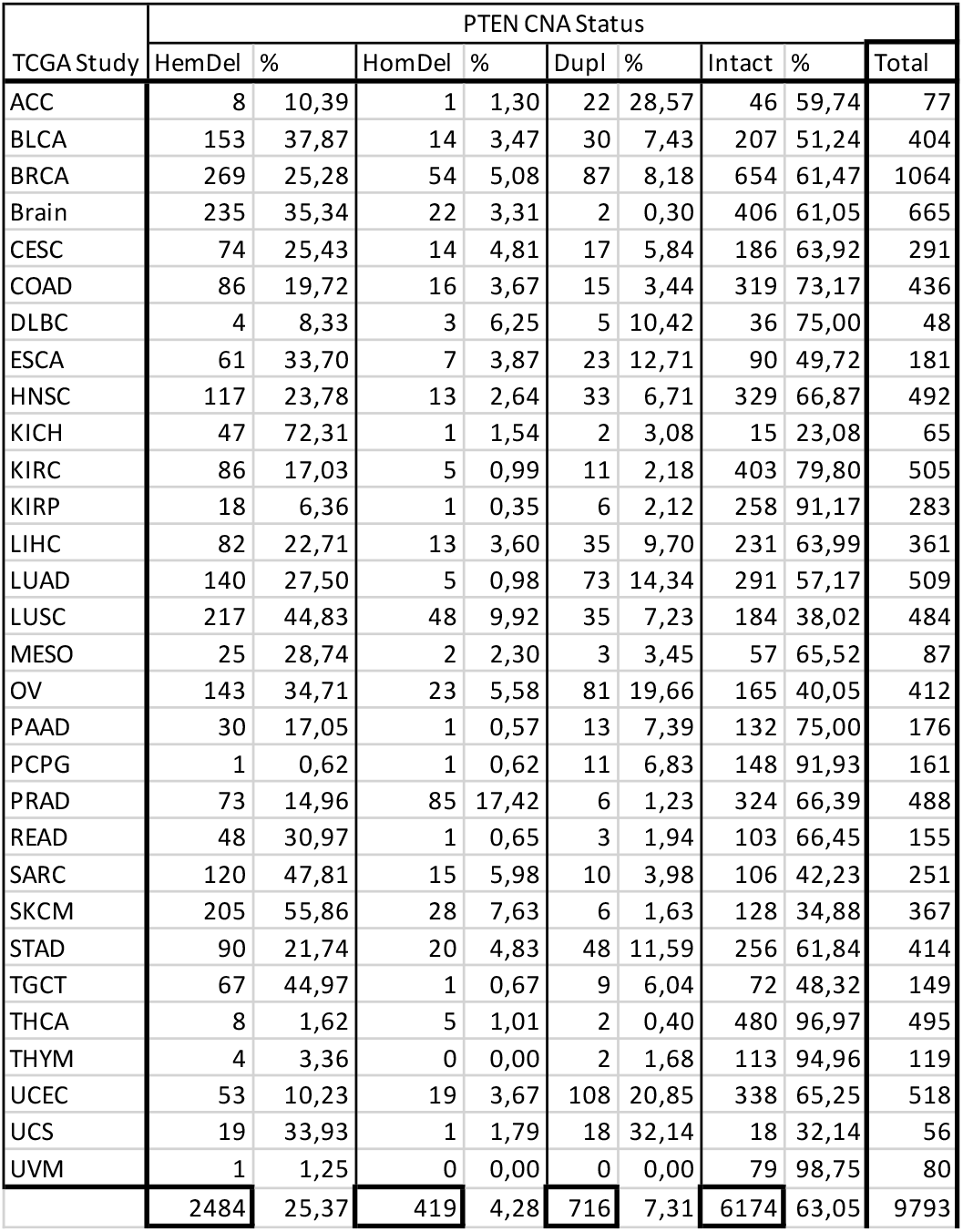
*PTEN* somatic copy number status (SCNA). HemDel: *PTEN* hemizygous deletion. HomDel: *PTEN* homozygous deletion. *Dupl* Duplication or gain across the Pan-cancer cohort of 30 tumors. ACC - Adrenocortical carcinoma, BLCA - Bladder Urothelial Carcinoma, BRCA - Breast invasive carcinoma, Brain (LGG - Brain Lower Grade Glioma combined with glioblastoma GBM), CESC - Cervical squamous cell carcinoma and endocervical adenocarcinoma, COAD - Colon adenocarcinoma, DLBC - Lymphoid Neoplasm Diffuse Large B - cell Lymphoma, ESCA - Esophageal carcinoma, HNSC - Head and Neck squamous cell carcinoma, KICH – kidney chromophobe, KIRC - Kidney renal clear cell carcinoma, KIRP - Kidney renal papillary cell carcinoma, LIHC - Liver hepatocellular carcinoma, LUAD - Lung adenocarcinoma, LUSC – Lung Squamous Squamous Cell Carcinoma, MESO - Mesothelioma, OV - Ovarian serous cystadenocarcinoma, PAAD - Pancreatic adenocarcinoma, PCPG - Pheochromocytoma and Paraganglioma, PRAD, Prostate Adenocarcinoma, READ - Rectum adenocarcinoma, SARC - Sarcoma, SKCM - Skin Cutaneous Melanoma, STAD - Stomach adenocarcinoma, TGCT - Testicular Germ Cell Tumors, THCA - Thyroid carcinoma, THYM - Thymoma, UCEC - Uterine Corpus Endometrial Carcinoma, UCS - Uterine Carcinosarcoma, UVM - Uveal Melanoma.

| TCGA Study | PTEN CNA Status |  |  |  |  |  |  |  | Total |
| --- | --- | --- | --- | --- | --- | --- | --- | --- | --- |
|  | HemDel | % | HomDel | % | Dupl | % | Intact | % |  |
| ACC | 8 | 10,39 | 1 | 1,30 | 22 | 28,57 | 46 | 59,74 | 77 |
| BLCA | 153 | 37,87 | 14 | 3,47 | 30 | 7,43 | 207 | 51,24 | 404 |
| BRCA | 269 | 25,28 | 54 | 5,08 | 87 | 8,18 | 654 | 61,47 | 1064 |
| Brain | 235 | 35,34 | 22 | 3,31 | 2 | 0,30 | 406 | 61,05 | 665 |
| CESC | 74 | 25,43 | 14 | 4,81 | 17 | 5,84 | 186 | 63,92 | 291 |
| COAD | 86 | 19,72 | 16 | 3,67 | 15 | 3,44 | 319 | 73,17 | 436 |
| DLBC | 4 | 8,33 | 3 | 6,25 | 5 | 10,42 | 36 | 75,00 | 48 |
| ESCA | 61 | 33,70 | 7 | 3,87 | 23 | 12,71 | 90 | 49,72 | 181 |
| HNSC | 117 | 23,78 | 13 | 2,64 | 33 | 6,71 | 329 | 66,87 | 492 |
| KICH | 47 | 72,31 | 1 | 1,54 | 2 | 3,08 | 15 | 23,08 | 65 |
| KIRC | 86 | 17,03 | 5 | 0,99 | 11 | 2,18 | 403 | 79,80 | 505 |
| KIRP | 18 | 6,36 | 1 | 0,35 | 6 | 2,12 | 258 | 91,17 | 283 |
| LIHC | 82 | 22,71 | 13 | 3,60 | 35 | 9,70 | 231 | 63,99 | 361 |
| LUAD | 140 | 27,50 | 5 | 0,98 | 73 | 14,34 | 291 | 57,17 | 509 |
| LUSC | 217 | 44,83 | 48 | 9,92 | 35 | 7,23 | 184 | 38,02 | 484 |
| MESO | 25 | 28,74 | 2 | 2,30 | 3 | 3,45 | 57 | 65,52 | 87 |
| OV | 143 | 34,71 | 23 | 5,58 | 81 | 19,66 | 165 | 40,05 | 412 |
| PAAD | 30 | 17,05 | 1 | 0,57 | 13 | 7,39 | 132 | 75,00 | 176 |
| PCPG | 1 | 0,62 | 1 | 0,62 | 11 | 6,83 | 148 | 91,93 | 161 |
| PRAD | 73 | 14,96 | 85 | 17,42 | 6 | 1,23 | 324 | 66,39 | 488 |
| READ | 48 | 30,97 | 1 | 0,65 | 3 | 1,94 | 103 | 66,45 | 155 |
| SARC | 120 | 47,81 | 15 | 5,98 | 10 | 3,98 | 106 | 42,23 | 251 |
| SKCM | 205 | 55,86 | 28 | 7,63 | 6 | 1,63 | 128 | 34,88 | 367 |
| STAD | 90 | 21,74 | 20 | 4,83 | 48 | 11,59 | 256 | 61,84 | 414 |
| TGCT | 67 | 44,97 | 1 | 0,67 | 9 | 6,04 | 72 | 48,32 | 149 |
| THCA | 8 | 1,62 | 5 | 1,01 | 2 | 0,40 | 480 | 96,97 | 495 |
| THYM | 4 | 3,36 | 0 | 0,00 | 2 | 1,68 | 113 | 94,96 | 119 |
| UCEC | 53 | 10,23 | 19 | 3,67 | 108 | 20,85 | 338 | 65,25 | 518 |
| UCS | 19 | 33,93 | 1 | 1,79 | 18 | 32,14 | 18 | 32,14 | 56 |
| UVM | 1 | 1,25 | 0 | 0,00 | 0 | 0,00 | 79 | 98,75 | 80 |
|  | 2484 | 25,37 | 419 | 4,28 | 716 | 7,31 | 6174 | 63,05 | 9793 |

### Hemizygous *PTEN* deletions (HemDel) are more common than homozygous deletions (HomDel) across the pan-cancer cohort

For most of the common solid tumors, hemizygous *PTEN* deletions were much more frequent than homozygous deletions (Figure 1). For example, melanoma (SKCM) has 205 (55.86%) hemizygous deletion compared to 28 (7.63%) homozygous loss. Breast cancer (BRCA) has a hemizygous deletion frequency of 269 (25.28%), but homozygous deletions are less common at 54 (5.08%) of tumors. There are interesting differences in lung subtypes, with squamous cell carcinoma (LUSC) having 217 (44.83%) hemizygous and 48 (9.92%) homozygous losses. However, adenocarcinoma (LUAD) had fewer hemizygous deletions, 140 (27.50%) but only 5 (0.98%) homozygous losses. For prostate cancer (PRAD), the rate of hemizygous loss was 73 (14.96%), slightly less than the homozygous loss rate of 85 (17.42%). The four tumor types (KIRP, THYM, THCA, and PCPG) with the lowest frequency of *PTEN* SCNA are characterized by a relatively more favorable outcome in the literature. Pheochromocytoma and paraganglioma (PCPG) are tumors of the autonomic nervous system. This tumor type has 11 (6.83%) *PTEN* duplication gains suggesting instability leading to *PTEN* acquisition rather than loss may be a feature of this tumor. Other tumors with more marked levels of *PTEN* SCNA gain are ovarian (OV) 81 (19.66%), esophageal carcinoma (ESCA) 23 (12.71%), lung adenocarcinoma (LUAD) 73 (14.34%), stomach adenocarcinoma (STAD) 48 (11.59%) and hepatocellular carcinoma of the liver (LIHC) 35 (9.70%).

**Figure 1.**
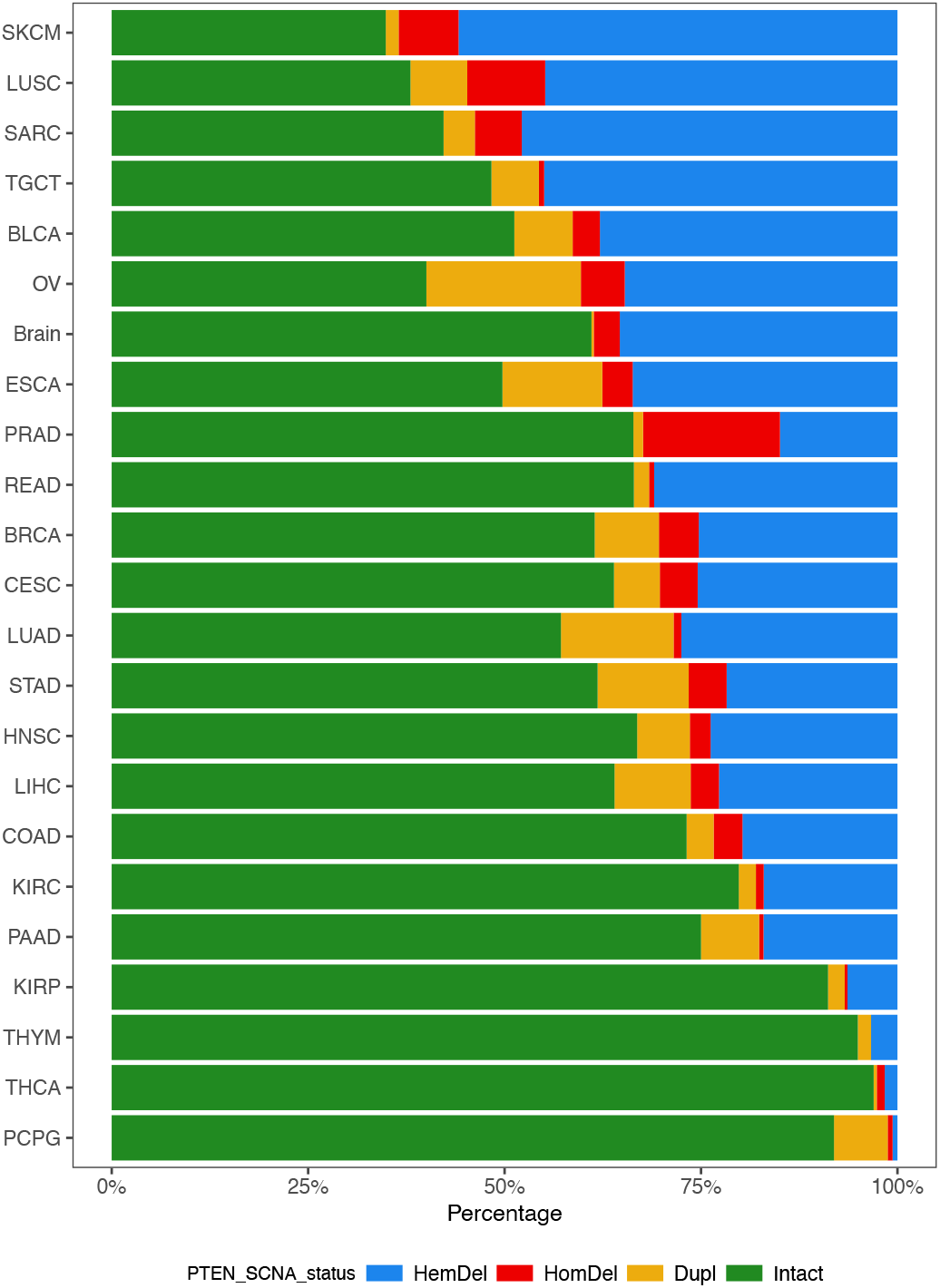
Relative percentages of *PTEN* Somatic Copy Number Alterations (*PTEN* SCNA) from the 23 most common solid tumors taken from the 30 tumor types analyzed in the Pan-cancer cohort (shown in Table 1). Hemizygous deletions (HemDel) = blue; Homozygous deletions (HomDel) = red, Intact (no *PTEN* SCNA) = green, and gain/duplication = brown. The tumors are ranked by their combined frequencies of PTEN deletion (HemDel + HomDel) with melanomas (SKCM) having the highest overall frequency of *PTEN* deletion (top) and Pheochromocytoma and Paraganglioma (PCPG) having the lowest (bottom). Duplication or *PTEN* gain (Dupl) was not included in the rank order determination. Plotted using *ggplot2* (https://ggplot2.tidyverse.org/).

### PTEN expression levels closely correlate to gene copy number

To determine how closely *PTEN* deletions and gains affect *PTEN* expression, we analyzed transcript counts across the pan-cancer cohort to estimate gene expression levels related to *PTEN* copy number (Figure 2a). We observed that tumors with *PTEN* HomDel had the lowest expression levels, with mean transcript counts that were one order of magnitude lower than the mean count observed in intact tumors (P<0.0001). The HemDel tumors showed transcript counts lower than intact (P<0.0001), and *PTEN* gains were associated with slightly higher expression levels than *PTEN* intact tumors (P<0.0001). We compared *PTEN* intact to HemDel and HomDel in the cohort’s 23 most common solid tumors (Figure 2b). The same reduced expression pattern is apparent in all tumor types, with *PTEN* HomDel having the lowest expression levels for most tumors. Similarly, *PTEN* HemDel exhibited intermediate expression levels between HomDel and *PTEN* intact tumors. These data agree with the general correlations observed between gene copy number and RNA expression levels using TCGA transcriptomic data reported recently^27^.

**Figure 2.**
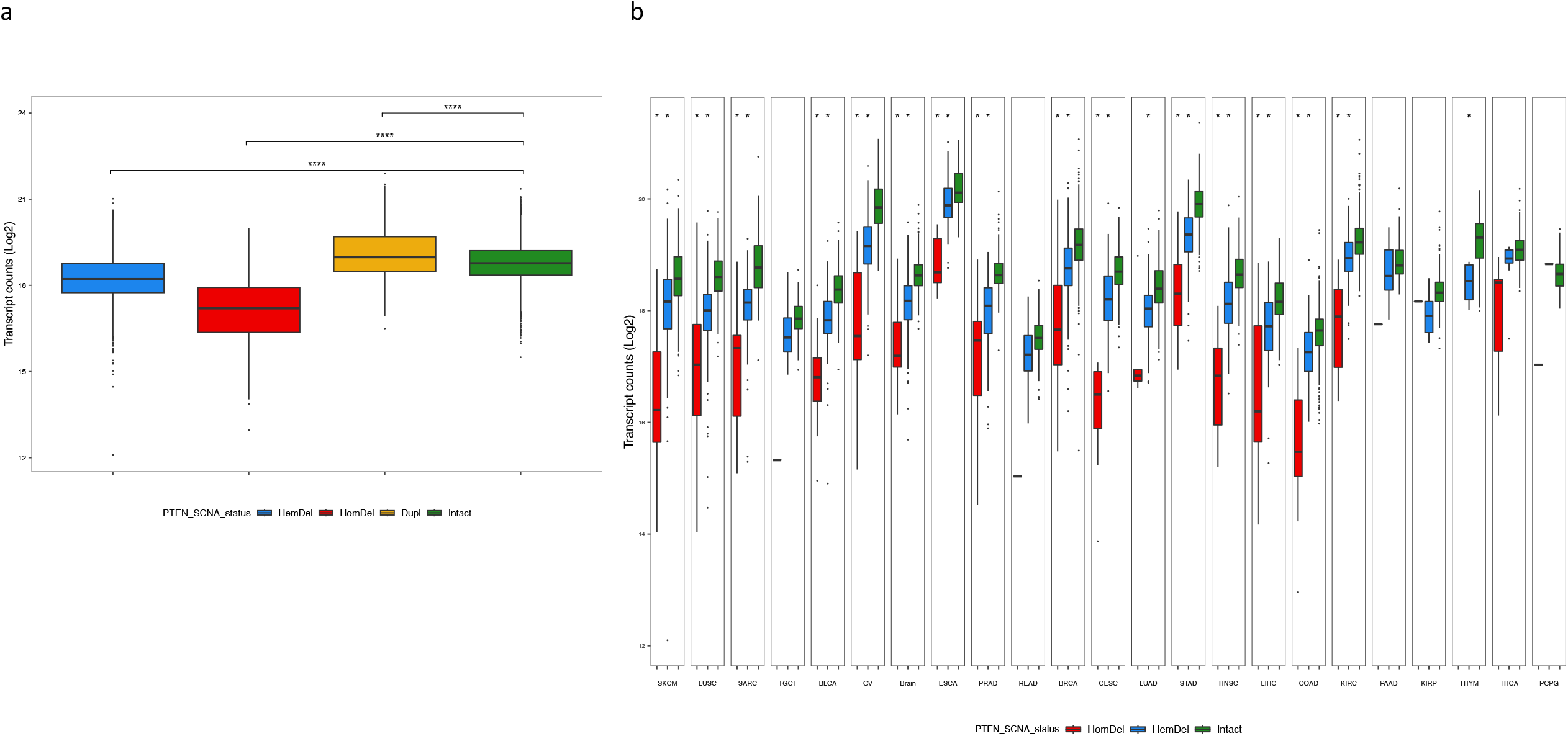
(**a**) Combined analysis of the association between *PTEN* SCNA status and differential gene expression of PTEN for all 30 tumors in the Pan-cancer cohort. Boxplots show the relationship between PTEN normalized transcript counts Log^2^ (Y-axis) versus *PTEN* SCNA status (HemDel, HomDel, Dupl and Intact). The mean transcript count for *PTEN* intact was 18.78, HomDel was the lowest expression level at 17.15, and HemDel showed reduced expression at 18.26, but Dupl was slightly higher than intact at 19.20. The mean transcript counts for all three classes of SCNA significantly deviated from intact (P<0.0001, t-test). (**b**) Association between *PTEN* SCNA status and differential gene expression of PTEN for the 23 most common solid tumors in the Pan-cancer cohort. Boxplots show the relationship between PTEN transcript counts Log^2^ (Y-axis) versus *PTEN* SCNA status HomDel (red), HemDel (blue), and Intact (green) for each tumor. For most of tumors, the mean transcript counts for both HomDel and HemDel deviated significantly from intact (P<0.05). Plotted using *ggplot2* (https://ggplot2.tidyverse.org/).

### Associations between hemizygous *PTEN* deletions and poor outcome

Analysis of the overall survival of the entire cohort stratified by *PTEN* deletion status showed that patients with tumors harboring *PTEN* HemDel had a similar poor outcome to those with HomDel (Figure 3a). We next performed outcome analysis comparing *PTEN* intact to HomDel and HemDel for four solid tumors (Figure 3b-e) to illustrate the diversity of effects of *PTEN* loss on survival. For colorectal carcinomas and head and neck cancers, the proportion of tumors with HomDel *PTEN* loss was relatively low, so these patients were not considered for comparative studies. In brain tumors, the *PTEN* intact have a steadily decreased survival probability, but both *PTEN* HomDel and HemDel were associated with a rapid decline in survival probabilities (Figure 3b). In cervical tumors, there is a lower survival probability for *PTEN* HemDel compared to HomDel and intact (Figure 3c). In stomach tumors (Figure 3d), HemDel was associated with intermediate survival risk. In head and neck squamous cell carcinoma (Figure 3e), the HemDel tumors were associated with reduced survival probabilities. *TP53* mutation was more common than expected for all HemDel tumors across the cohort (Supplementary Table 1). Further analysis considering the combined effects of *TP53* mutation with *PTEN* deletion status was performed (Supplementary Table 2). Similar hazard ratios were observed for *TP53* monoallelic loss and *PTEN* HemDel, and there was no obvious additive effect when both mutations occurred together (Supplementary Table 2a). HNSCC tumors also showed similar hazard ratios for *PTEN* HemDel and *TP53* monoallelic mutation without apparent synergy when both mutations were present (Supplementary Table 2b). These data suggest that one copy loss of either *PTEN* or *TP53* results in equivalent effects on cancer outcome.

**Figure 3.**
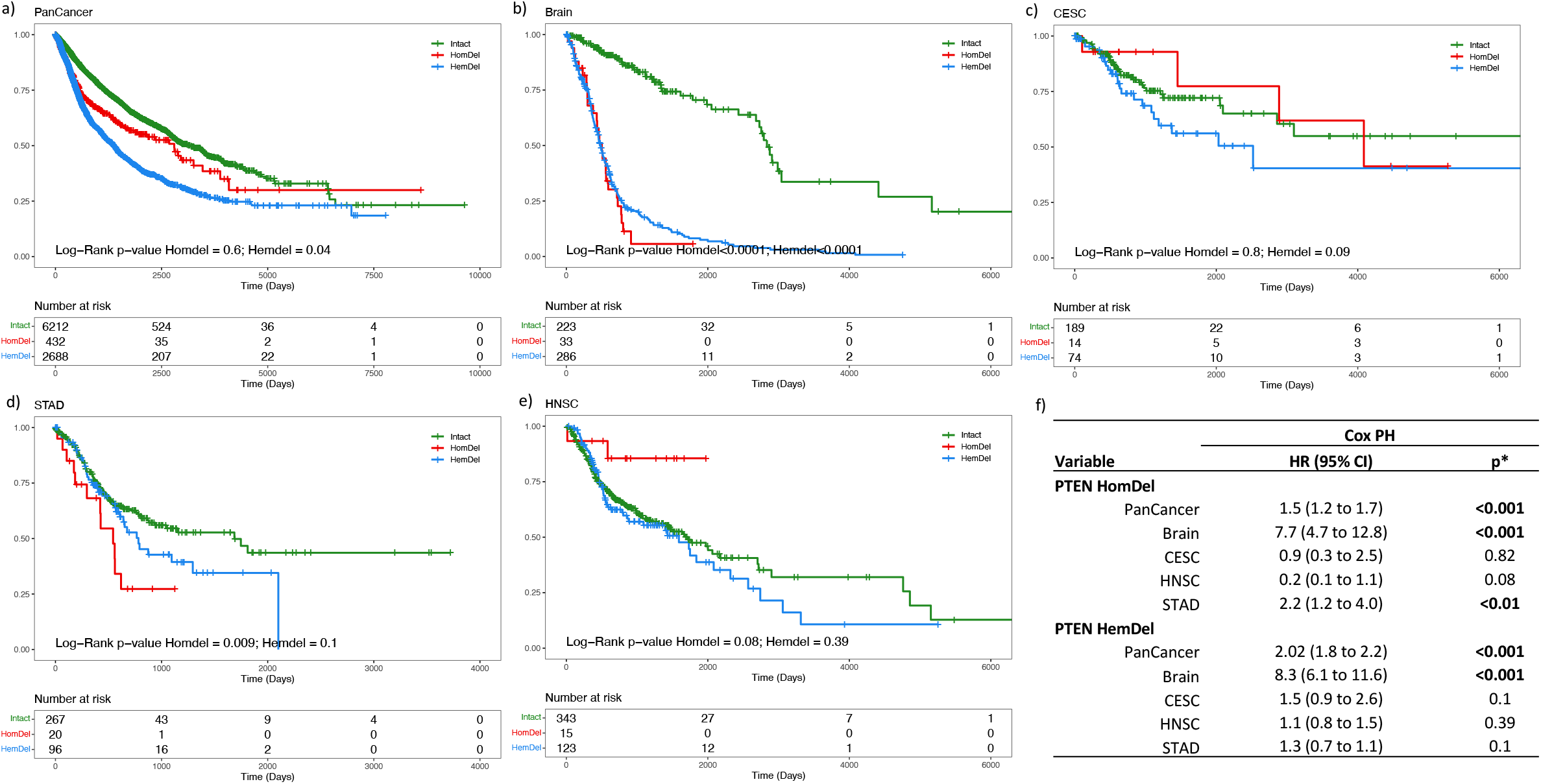
Kaplan-Meier curves illustrate overall survival for both PTEN intact and PTEN deletions groups (HemDel and HomDel). Boxplots **(a)** Pan-cancer cohort shows a lower survival probability for PTEN HemDel compared to HomDel and intact (P<0.0001). **(b)** Brain tumors (GBM + LGG) show that both HomDel and HemDel are equally associated with earlier events (P<0.0001). **(c)** Cervical tumors show a lower survival probability for PTEN HemDel compared to HomDel and intact. **(d)** Head and neck cancer. **(e)** Stomach tumors. **(f)** Cox Model analysis for *PTEN* intact and *PTEN* deletions groups. Plotted using *Survminer* (https://rpkgs.datanovia.com/survminer/index.html).

### Relationships between hemizygous *PTEN* deletions and genomic instability

To understand the impact of *PTEN* loss on genomic features of cancer progression, we analyzed the total number of mutations, percent genome altered by losses and gains, intratumor heterogeneity, aneuploidy, and homologous repair defects across the pan-cancer cohort (Figure 4a-d). Our combined analysis revealed that cancers harboring *PTEN* HemDel had the highest levels of nonsilent mutations, intratumor heterogeneity, and percent genome altered. Moreover, PTEN HomDel and HemDel presented higher levels of homologous repair defect compared with *PTEN* intact, but no difference was found between the two deletion types. When stratified by tumor type, many of the tumors harboring *PTEN* deletions had increased levels of nonsilent mutations, intratumor heterogeneity, and homologous repair defects (Supplementary Figure 2a-c).

**Figure 4.**
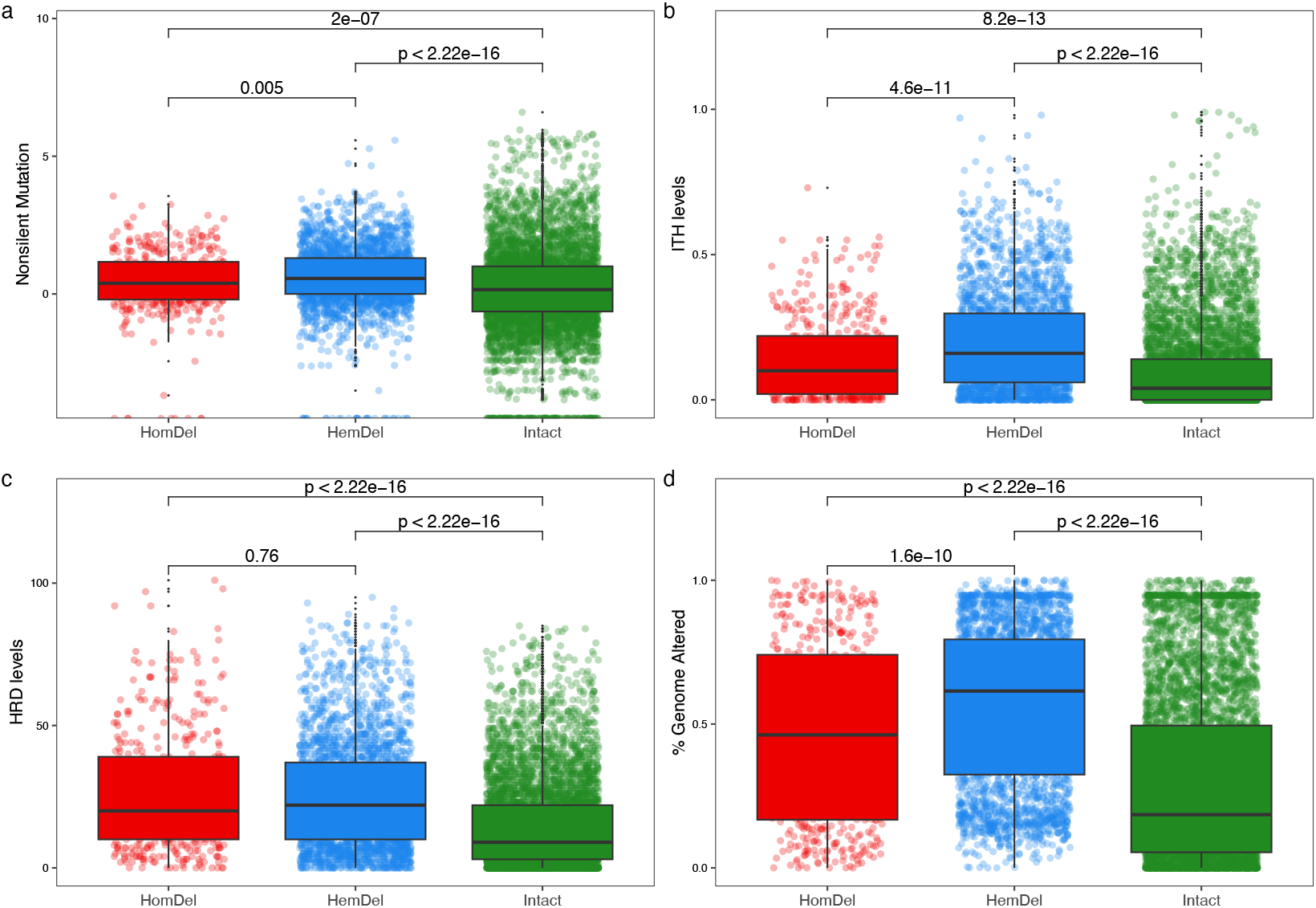
*PTEN* deletions are associated with (**a**) increased mutation levels, (**b**) intra-tumor heterogeneity (ITH), (**c**) homologous repair defects (HRD), and (**d**) percent genome alteration. Hemizygous deletions of *PTEN* are associated with increased levels of genomic alterations compared with homozygous deletions. Intact *PTEN* showed the lowest levels of genomic alterations. Plotted using *ggplot2* (https://ggplot2.tidyverse.org/).

To compare the percentage of genome altered, we found that 65% (13/20) of the investigated tumor types presented elevated genomic alterations when harboring *PTEN* HemDel compared to *PTEN* Intact tumors (Supplementary Figure 2d). Pan-cancer analysis showed aneuploidy levels were significantly elevated in HemDel compared to HomDel and both deletion types showed significantly increased aneuploidy compared to all intact tumors (Supplementary Figure 3a). Similar intratumor heterogeneity and aneuploidy patterns were found in Dupl cases (Supplementary Figure 4a-b). Analysis of aneuploidy in a subset of twenty solid tumors showed a general increase in aneuploidy associated with PTEN deletion. Hemizygous losses in bladder, cervix, colon, head and neck, liver, ovarian, pancreatic, and stomach cancer promoted increased levels of aneuploidy (Supplementary Figure 3b). Collectively, these findings led us to hypothesize that *PTEN* HemDel may be associated with an altered transcriptome, potentially driving more genomic alterations.

### Transcriptomic changes associated with *PTEN* loss activate immune pathways

We next investigated the transcriptome of seven representative solid tumors with increased frequencies of *PTEN* deletions and presented more diversity in genomic changes. Differential gene expression analyses showed prostate tumors with *PTEN* HemDel presented 2076 upregulated and 2966 downregulated genes compared to *PTEN* Intact tumors (Supplementary Table 3). Prostate tumors with HemDel of *PTEN* presented several enriched pathways compared with *PTEN* intact, including microtubule-based movement and chromosome segregation (Supplementary Figure 5). *PTEN* HemDel was also associated with distinct enrichment of immune pathways across the cancer types. In prostate tumors with *PTEN* HomDel the high number of DEGs was linked to cell cycle, DNA repair, and vascular process pathways (Supplementary Figure 6). Interestingly, for HomDel in prostate cancer, the genes closely flanking *PTEN*, such as *ATAD1, RNLS* and *KLLN*, are concurrently downregulated with the *PTEN* gene consistent with the large genomic deletions of chromosome 10 previously observed in prostate cancer by FISH ^28^ (see the vertical bar in Supplementary Figure 6).

Brain tumors presented the most distinct pathway enrichment results, including the upregulation of several immune pathways when *PTEN* was deleted. We found 3562 up- and 10902 downregulated genes in these tumors when compared with *PTEN* HemDel vs. *PTEN* intact (Supplementary Table 3). GO-enriched analysis showed that several immune-related pathways were upregulated for *PTEN* HemDel tumors, such as granulocyte migration and chemotaxis, neutrophil migration, acute inflammatory response, humoral immune homeostasis, and many others (Supplementary Figure 7). Similarly, when we compared the transcriptome of tumors with *PTEN* HomDel vs. those with PTEN intact, we observed 4106 up- and 7332 downregulated genes (Supplementary Table 3, Supplementary Figure 8). We also observed that several immune-related pathways were significantly upregulated in brain tumors harboring *PTEN* HomDel, namely myeloid leucocyte migration, humoral immune response, leucocyte chemotaxis, and acute inflammatory response (Supplementary Figure 8). In summary, most investigated tumors had immune-related pathways up- or downregulated when *PTEN* was lost. The RNA-sequencing expression level of immunomodulatory genes like CD274 (PD-L1, Programmed Cell Death), CTLA4 (Cytotoxic T-Lymphocyte Associated Protein 4), IDO1 (Indoleamine 2,3-Dioxygenase 1), and LAG3 (Lymphocyte Activating 3) are altered in breast, glioblastoma, head and neck, liver, ovarian, prostate, sarcoma, stomach, and uterine tumors (Supplementary Figure 9). These findings led us to hypothesize whether *PTEN* genomic deletions may be associated with an altered tumor microenvironment, as determined from higher or lower abundance of immune cells surrounding cancerous lesions.

Stomach tumors harboring *PTEN* HemDel show downregulated immune pathways, such as adaptive immune response, regulation of lymphocyte activation, T cell activation, positive regulation of immune response, leukocyte-mediated immunity, lymphocyte proliferation, and several other pathways (Supplementary Figure 10). Stomach tumors with *PTEN* HemDel also had downregulation of Indoleamine 2,3-Dioxygenase 1 (*IDO1*) gene (Supplementary Figure 10). In contrast, activation of immune pathways was not apparent in stomach tumors harboring *PTEN* HomDel, but the sample size (N=20) for homozygous loss tumors was reduced compared to HemDel tumors (N=90) (Supplementary Figure 11, Supplementary Table 3). For head and neck tumors, we also observed that tumors with *PTEN* HemDel had downregulated immune-related pathways, including adaptive immune response, activation of T-cell receptor, regulation of lymphocyte-mediated immunity, regulation of leukocyte-mediated immunity, and regulation of immune effector process (Figure 5a-c). However, these results were not observed for the comparison between *PTEN* HomDel and *PTEN* intact, in which the sample size from homozygous loss tumors was limited (N=15). However, the linked passenger genes *MINPP1* and *KLLN* from chromosome 10 showed concurrent reduced expression with *PTEN* (Supplementary Figure 12). Lastly, for colon tumors, we found that tumors with *PTEN* HemDel had downregulated immune-related pathways, including response to interferon-gamma, lymphocyte-mediated immunity, adaptive immune response, and natural killer cell-mediated immunity (Supplementary Figure 13). However, effects on immune pathways were not observed for the comparison between *PTEN* HomDel and *PTEN* intact. As with other homozygous losses, there was reduced expression of a group of *PTEN*-linked passenger genes mapping to chromosome 10, including *ALDH18A1, DNMBP, KLLN, WAPL, MINPP1*, and *ATAD1* (see the vertical bar in Supplementary Figure 14). Collectively, the pathway analysis results draw attention to the role hemizygous *PTEN* loss may play in the anticancer immune response.

**Figure 5.**
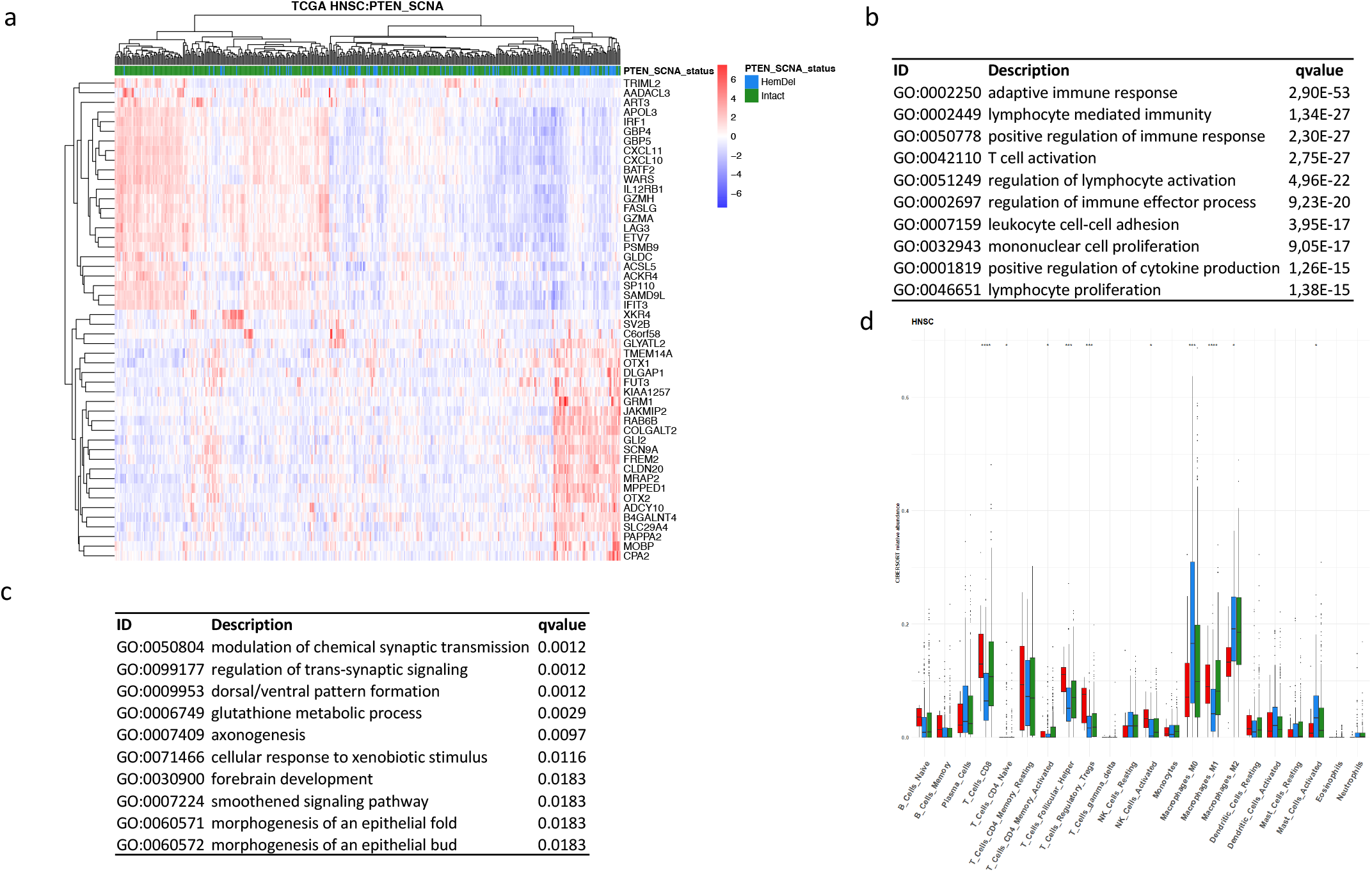
*PTEN* HemDel in head and neck squamous cell carcinoma (HNSC) shows distinct transcriptome activity. (**a**) Transcriptome heatmap exhibiting clustering of top 50 DEGs (Log2Fold change >0.58). The color scale in the heatmap represents the Z-score of the normalized read counts for each gene, where the red scale color indicates upregulated, and blue low expressed genes. Enrichment analyses were performed using *clusterProfiler* and *p*-adjusted value = 0.05 as the cutoff. (**b**) Enriched GO (BP) pathways of the upregulated genes. (**c**) Enriched GO (BP) pathways of the downregulated genes. The tables shown in (b) and (c) indicate the top 10 most significant Gene Ontology (BP) terms. (**d**) CIBERSORT□derived immune□cell abundance of 22 distinct cell subsets based on *PTEN* status and head and neck tumors. Y□axis shows CIBERSORT relative scores. *PTEN* HemDel in head and neck tumors was linked to higher M0 and M2 macrophages, mast cells, and lower CD4 T, CD8 T, T regs, and NK cells. *: *p*-value < 0.05; **: *p*-value < 0.01, by Mann-Whitney test. n=492. DC, dendritic cell; NK, natural□killer cell; PCa, prostate cancer; Treg, regulatory T cell; M0, M0 macrophage; M1, M1 macrophage; M2, M2 macrophage. Plotted using *pheatmap* (https://rdocumentation.org/packages/pheatmap/versions/1.0.12).

### *PTEN* deletions are associated with an altered tumor immune microenvironment

The results from the tumor microenvironment comparison of *PTEN* HomDel, HemDel, and intact tumors for all 22 immune cell types from brain, prostate, head and neck, and stomach tumors were investigated. *PTEN* HemDel in brain tumors was linked to higher CD4 T memory cells and M2 macrophage abundance and lower M1 macrophages, monocytes, and CD8 T cells (Supplementary Figure 15a). Similar results were seen in head and neck tumors, where *PTEN* HemDel showed higher M0 and M2 macrophages and mast cells and lower CD4 T-cell, CD8+ T-cell, T-regulatory cells, and NK cells (Figure 5d). While in prostate tumors with *PTEN* HemDel, we observed an increase in M1 macrophages and a decrease in M0 macrophages and monocytes abundance (Supplementary Figure 15b). Stomach tumors with *PTEN* HemDel had an increased plasma and M0 macrophages density and decreased density of CD8 T-cells (Supplementary Figure 15c). Cervical tumors with *PTEN* HemDel were linked to higher M0 macrophages and an abundance of activated mast cells (Supplementary Figure 15d). Similar results were seen in colorectal tumors, where *PTEN* HemDel showed a higher M0 macrophage density and lower CD8 T-cell abundance (Supplementary Figure 15e).

## DISCUSSION

The presence of the PTEN protein is essential for controlling the tumor promoting activities of the PI3K/AKT pathway ^2^. Deregulation of this signaling axis confers a strong selective advantage to tumors, so that loss of *PTEN* control by inactivating mutation or deletion is one of the most common somatic events in human cancer ^6^. PTEN mutation is significantly related to advanced disease, chemotherapy resistance, and poor survival in patients with head and neck cancers, breast, melanoma, colorectal, esophageal, and prostate ^19,29–34^ so that its role as a predictive and prognostic cancer biomarker is increasingly gaining prominence ^35–37^. Moreover, there is increasing awareness of the role PTEN deficiency plays in shaping the immune landscape in different cancers ^3,4^.

Various murine models have shown that the tumor-promoting abilities of loss of the *Pten* gene are strongly dose-dependent ^38^. In some models, the *Pten* gene loss has been found to exhibit haploinsufficiency with reduced levels of the normal Pten protein resulting in a diversity of tumors in mice ^2^. Thus, in contrast to the classical definition of a tumor suppressor gene that requires complete loss of both copies of the gene ^39^, it appears that reduced amounts of PTEN protein caused by either monoallelic loss or by one inactivating mutation is sufficient to cause cancer (reviewed in ^19^).

Partial loss of function caused by *PTEN* hypomorphic mutations has been observed amongst PTEN hamartoma tumor syndrome (PHTS) patients ^40^. A lower cancer risk and possibly a higher likelihood of neurodevelopmental phenotypes, specifically autism spectrum disorder (ASD), was associated with *PTEN* hypomorphic mutations.

A recent review across all solid tumors showed that hemizygous *PTEN* loss due to monoallelic deletion or point mutation occurred at a much higher frequency than biallelic loss ^15^. Our *PTEN* analysis of TCGA data also shows that most tumors have a much greater frequency of HemDel than HomDel and that hemizygous loss is less favorable than homozygous loss for some tumor types. These data agree with previous genetic and proteomic studies of PI3K/AKT/mTOR axis mutations using TCGA data that showed outcomes associated with partial loss of *PTEN* were less favorable than outcomes for complete loss of the gene ^18^. Our analysis of *PTEN* RNA transcript levels across the TCGA cohort shows good agreement with gene copy number, as reported recently by others ^27^, so the HemDel losses we analyzed in this study would be expected to reduce RNA expression levels. Moreover, since one somatic *PTEN* deletion or point mutations leads to a reduction in RNA levels, then partial PTEN protein expression is expected in HemDel tumors.

*PTEN* deletions and protein loss are known to affect DNA damage responses and disrupt mitotic spindle architecture resulting in accumulation of mutations in other genes and increased chromosome instability in both murine models and human cancers ^41^. As expected, our TCGA analysis showed both hemi- and homozygous losses increased genomic instability features in many tumors. However, HemDel tumors had significantly higher rates of non-silent mutations, intratumor heterogeneity, percentage genome altered, and aneuploidy compared to HomDel tumors. This diversity of genetic alteration is likely to facilitate genomic evolution and increase cancer progression, treatment resistance, and poor outcome ^42^. Collectively, these data suggest that the elevated levels of genomic instability and somatic mutations in the tumors with hemizygous loss could provide a selective advantage for the acquisition of other driver genes and the less favorable outcomes observed across the TCGA cohort.

PTEN loss may be sufficient to cause tumorigenesis in some tissues but not in others. A recent TCGA analysis of collaborating somatic mutations in *PTEN*-defective solid tumors identified cancer genes such as *TP53, APC, TTN, MUC16, PIK3CA, BRAF, CDH1*, and *KMT2D*, irrespective of the tumor type ^37^. When surveying the outcome in different cancers across the TCGA cohort, we found the consequences of HemDel loss were more pronounced in some tumor types, suggesting that other cellular contexts or unknown tumor-specific molecular features may contribute to progression when *PTEN* is partially deficient.

The most obvious effects on outcome for HemDel *PTEN* losses were observed in head and neck squamous cell carcinomas (HNSCC) harboring a mono-allelic somatic loss-of-function mutation in *TP53*. For this type, the hemizygous loss was associated with a less favorable overall survival with a significant HR of 1.67 (P=0.023) (Figure 3, Supplementary table 2). *PTEN* loss is associated with poor outcomes in HNSCC ^30^. HNSCC also has a high frequency of *TP53* somatic mutations ^43^. We used cBioPortal to investigate the co-occurrence frequencies of *TP53* mutations with *PTEN* intact, HemDel, and HomDel in HNSCC. Interestingly, *PTEN* HemDel was significantly (Supplementary table 1, P<0.001) associated with *TP53* mutation, and *PTEN* HomDel was under-represented in HNSCC with mutated *TP53* (data not shown). These data suggest that there may be preferential selection for one copy of a *PTEN* gene in HNSCC so that reduced levels of the functional PTEN protein allow tumor cells to bypass the *TP53*-induced cell senescence pathway ^44^. These observations in HNSCC are in keeping with the *PTEN* literature that suggests that single-copy inactivation of *PTEN* may actually be selected for during the progression of some tumor types since bi-allelic inactivation of *PTEN* has been shown to lead to senescence or cell death when *TP53* is mutated ^5^. Recent functional studies of the relationship between *TP53* mutation and *PTEN* loss in HNSCC indicate that loss of regulation of both pathways could influence radio- and chemosensitivity ^45^. These findings also agree with recent studies showing that *PTEN* loss mediates resistance to cetuximab in HNSCC, indicating the need for more treatment response *PTEN* predictive biomarker studies that also examines the role of HemDel events ^46^.

Mouse models of HNSCC draw attention to the role of *Pten* in the evasion of cellular senescence and activation of cancer-related inflammation ^47^. Our gene set enrichment analysis for HNSCC indicates that various pathways involved with immune activation are downregulated in *PTEN* HemDel tumors. Pathways indicated in our analysis were antigen presentation, interferon, interleukins, toll-like, neutrophil, B cell, and inflammasome signaling. More marked alterations to the immune response in HemDel HNSCC were also evident in our CIBERSORT analysis. There were decreased CD8 T cells and alterations to tumor-associated macrophages (TAMs) consistent with the immune tumor microenvironment being less favorable in hemizygous HNSCC. These findings are consistent with the emerging recognition that *PTEN* loss could be a useful predictive immune biomarker in HNSCC ^48^.

Biomarker analysis of colorectal cancer showed weak expression of the PTEN protein in primary tumors with metastasis^33^. Our study found that *PTEN* HemDel colorectal tumors had downregulated immune-related pathways. Functional activities associated with immune response included regulation of granulocyte-macrophage colony-stimulating factor production, neutrophil migration, and regulation of leukocyte-mediated cytotoxicity. These findings are consistent with recent studies that showed that *PTEN* mutation was associated with microsatellite instability subtypes and tumor mutational burden in colon cancer ^49^. Both *in vitro* studies ^50^ and recent analyses of colon tumors ^51^ suggest that *PTEN* loss in colorectal carcinoma may activate PI3K and upregulate *PD-L1* expression. These data are in keeping with other studies indicating that the PTEN/ PI3K axis may promote immune escape through regulating *PDL1/PD1* expression ^3,52^.

Brain tumors also presented distinct pathway enrichment findings, including the upregulation of several immune pathways for *PTEN* HemDel and HomDel losses. These data are consistent with a recent study showing *PTEN* loss in glioblastoma was associated with immunosuppressive expression signatures and failed response to anti-PD-1 immunotherapy ^53^.

Our TCGA analysis of prostate tumors with *PTEN* HemDel identified elevated levels of genomic instability and increased M1 macrophages but decreased abundances of M0 macrophages and monocytes. Pathway analysis showed enrichment for response to type I-interferon, consistent with other recent findings showing that *PTEN* loss leads to an immunosuppressive microenvironment in prostate cancer (PCa) ^54^. These conclusions also agree with our previous study that showed increased IDO1 and Tregs in *PTEN*-deficient prostate cancer tumor microenvironment ^55^. Interestingly, analysis of PCa showed that the genes flanking *PTEN*, such as *ATAD1, RNLS*, and *KLLN* are concurrently down-regulated when the *PTEN* gene is homozygously deleted. Similar observations were made in both colon and head and neck tumors with homozygous deletions. These data suggest that large somatic deletions of chromosome 10 may often encompass closely linked genes whose expression changes could contribute to clinical phenotypes ^36^.

Unfortunately, most previous *PTEN* cancer biomarker studies have not systematically analyzed mutations or deletions using molecular genetic methods that can distinguish between HemDel and HomDel mutations. Similarly, loss of the protein using immunohistochemistry as an alternative way of determining PTEN status has also not been studied using uniform, reproducible methods that can detect subtle changes in protein levels ^15,17,25,26^. The end result has been that majority of published *PTEN* cancer biomarker studies have depended on a variety of simple two-way classification assays (*PTEN* loss vs. intact) that ignore the distinction between hemi- and homozygous losses. Our study also has some limitations that need to be addressed in future *PTEN* analyses of the highlighted tumors. Our work was based entirely on *in silico* analysis of copy number data derived TCGA, and all the pathway and immune parameters were inferred from the associated transcriptome data. The analysis presented here cannot fully capture the impact of *PTEN* mutations based on the limitations of the TCGA dataset, which lacks prognostic or treatment information. Also, our expression analyses associated with *PTEN* status were, like all *in silico* TCGA studies, obtained from the bulk tumors containing heterogeneous tumor cells intermingled with a small subset of stromal cells and varying quantities of immune cells. Lastly, our analysis regarding *PTEN* status did not include *PTEN* inversions, fusions, inactivating somatic mutations, promoter methylations, and large-scale structural variants, which may be a source of bias in our evaluation of the impact of HemDel and HomDel effects.

Collectively, our findings based on TCGA, together with the extensive murine *Pten* studies by others ^8,9^, suggest that loss of one copy of *PTEN* by mutation or deletion may be sufficient to promote tumorigenesis and anti-cancer immune evasion. However, for most human cancers, there have been limited biomarker studies on how hemizygous *PTEN* deletion and partial protein loss may influences outcome and treatment response.

## METHODS

### Data download and processing

We initially downloaded raw and level 3 PanCancer normalized RNAseq, array-CGH, and SNV from 10,713 samples derived from 31 tumor types from The Cancer Genome Atlas (TCGA) cohort (www.gdc.cancer.gov). These initial cases were used to determine outcome, genomic alterations, and relative immune abundances. For the differential gene expression analysis (DEG), TCGA biolinks had 9793 cases were available from 30 primary tumors as detailed in Table 1.

Raw RNAseq experiments were performed through Illumina HiSeq 2000 RNA Sequencing platform. Copy number data were obtained from array-CGH experiments performed with the Affymetrix Genome-Wide Human SNP Array 6.0. DNA sequencing was performed in Illumina Genome Analyzer DNA sequencing. Clinical data for progression-free intervals and overall survival were obtained for all patients from the TCGA cohort. All analyses were conducted with data derived from primary tumors.

### Defining *PTEN* deletions

*PTEN* deletions were characterized by using GISTIC-derived categorical SCNA. In brief, GISTIC identifies a cutoff for copy number loss or gain using the log2 continuous data from each individual segments, such as genes and other regions ^56^. These analyses were performed by the TCGA group for the PanCancer analysis. We obtained, for our study, the Level 3 copy number calls for *PTEN* gene resulting from whole genome detection of copy number changes or point mutations inactivating either one or both copies of the gene ^20^. The calls were defined as HomDel (two copies with loss/mutation), HemDel (one copy loss or mutation), intact *PTEN* (both copies present without mutation), and duplications of the *PTEN* gene ^57^. *PTEN* duplications consisted of the presence of gain of an extra copy of the entire gene or regions at 10q23.31 that included *PTEN*. Our initial analysis of *PTEN* somatic mutations showed that in keeping with the literature ^15^, this class of inactivation of the gene was infrequent in the TCGA cohort (Supplementary Figure 1). We, therefore, choose to focus our analysis solely on SCNA affecting *PTEN*.

### Gene expression and enrichment analyses

Analysis of gene expression was performed on the RNAseq level. Raw read counts for TCGA-tumors were downloaded using FANTOM-CAT/recount2 (https://jhubiostatistics.shinyapps.io/recount/) ^58,59^. The subset of samples was compiled using each patient’s clinical information, and only primary tumors were used in the analysis. Study groups were divided based on *PTEN* status, HomDel (two copies with loss/mutation), HemDel (one copy loss or mutation), and Intact *PTEN* (both copies present without mutation). Differential expression (DE) analysis was carried out using *DESeq2* (v 1.32.0) 59,60, and *PTEN* status was used as the design factor. For enrichment analysis of DE genes, we used the clusterProfile package (v 3.18.1) 61,62. We used a P-adjusted value < 0.05 as the cutoff for the enrichment results. All analyses were conducted in RStudio software (R Foundation for Statistical Computing, R v4.1.2) and all results were plotted by *pheatmap* (v 1.0.12) and *ggplot2* ^64^. All results are displayed with *PTEN* Intact as a reference for statistical analysis.

### Immunogenic and genomic effects of PTEN deficiency

To determine the association between *PTEN* inactivation and acquired DNA changes in tumors, we investigated the presence of genome doublings, homologous recombination defects (HRD), aneuploidy levels, tumor mutational burden, and intratumoral heterogeneity. These data were downloaded from the PanCanAtlas database for The Immune Landscape of Cancer from TCGA (https://gdc.cancer.gov/about-data/publications/panimmune) ^65^.Genome doubling status was determined through ABSOLUTE algorithm ^66^, which measures tumor ploidy and purity based on copy number and mutational signatures. Ploidy levels were determined by the quantity of DNA that each cancer cell presents after undergoing several numerical and structural chromosome aberrations. Clonality calls were employed to determine intratumor heterogeneity scores, which determine tumor copy number and point mutations as aggregates of clonal and subclonal components having varying ploidy levels. Tumor purity is a measurement that considers the fractions of normal cells within the bulk mass of tumor cells. Aneuploidy levels were calculated based on the total length of gained and deleted chromosome arms divided by the genome size ^67^. Similarly, HRDs were calculated based on the copy number calls in with large (>15 Mb) non-arm-level chromosome imbalances with loss of heterozygosity, breaks between genes larger than >10 Mb, and subtelomeric regions harboring allelic imbalances. Groups with distinct *PTEN* inactivation statuses were compared by employing the Kruskal-Wallis test in R.

### Prognostic effect of PTEN-deficiency

To determine if *PTEN* HemDel and HomDel status has a significant impact in predicting recurrence and death events, we performed survival analysis using *Survival* package in R. Log-rank tests and Kaplan Meier curves were generated through the *Survival* and *Survminer* packages in R. In addition, Cox Regression univariate models were obtained for all tumors, grouped and separately by tumor type.

### *In silico* analysis of immune abundance in tumor types

To estimate the immune cell landscape in each tumor, we used the CIBERSORT deconvolution method (https://cibersort.stanford.edu/), which is an established tool to determine the abundance of immune cells using whole□transcriptome data ^68^. We separately imputed normalized gene expression data of brain, prostate, head and neck, stomach, cervical and colorectal tumors from the TCGA cohort. All results are displayed with *PTEN* Intact as a reference.

## DECLARATIONS

### Authors’ contributions

All authors contributed equally to the development of this study. All authors read and approved the final manuscript.

### Availability of data and materials

All data used in this analysis can be found at the GDC data portal. The code and processed data were provided as supplementary materials.

### Ethics approval and consent to participate

Research Ethics Committee of the Clinical Hospital of the Faculty of Medicine of Ribeirão Preto, number 3.088.034.

## REFERENCES

1. Liu, T., Wang, Y., Wang, Y. & Chan, A. M. Multifaceted regulation of PTEN subcellular distributions and biological functions. Cancers 11, (2019).

2. Chalhoub, N. & Baker, S. J. PTEN and the PI3-kinase pathway in cancer. Annual Review of Pathology 4, 127–150 (2009).

3. Vidotto, T. et al. Emerging role of PTEN loss in evasion of the immune response to tumours. British Journal of Cancer 122, 1732–1743 (2020).

4. Bezzi, M. et al. Diverse genetic-driven immune landscapes dictate tumor progression through distinct mechanisms. Nature Medicine 24, 165–175 (2018).

5. Lee, Y., Chen, M. & Pandolfi, P. P. The functions and regulation of the PTEN tumour suppressor: new modes and prospects. Nature Reviews Molecular Cell Biology 19, 547–562 (2018).

6. The ICGC/TCGA Pan-Cancer Analysis of Whole Genomes Consortium. Pan-cancer analysis of whole genomes. Nature 578, 82–93 (2020).

7. Bignell, G. R. et al. Signatures of mutation and selection in the cancer genome. Nature 463, 893–898 (2010).

8. Wang, S. et al. Prostate-specific deletion of the murine Pten tumor suppressor gene leads to metastatic prostate cancer. Cancer Cell 4, 209–221 (2003).

9. Kwabi-Addo, B. et al. Haploinsufficiency of the Pten tumor suppressor gene promotes prostate cancer progression. Proceedings of the National Academy of Sciences of the United States of America 98, 11563–11568 (2001).

10. Kwon, C. H. et al. Pten haploinsufficiency accelerates formation of high-grade astrocytomas. Cancer Research 68, 3286–3294 (2008).

11. Alimonti, A. et al. Subtle variations in Pten dose determine cancer susceptibility. Nature Genetics 42, 454–458 (2010).

12. Shen-Li, H., Koujak, S., Szablocs, M. & Parsons, R. Reduction of Pten dose leads to neoplastic development in multiple organs of PtenshRNA mice. Cancer Biology and Therapy 10, 1194–1200 (2010).

13. Carracedo, A., Alimonti, A. & Pandolfi, P. P. PTEN level in tumor suppression: How much is too little? Cancer Research 71, 629–633 (2011).

14. Masson, G. R. & Williams, R. L. Structural mechanisms of PTEN regulation. Cold Spring Harbor Perspectives in Medicine 10, (2020).

15. Álvarez-Garcia, V., Tawil, Y., Wise, H. M. & Leslie, N. R. Mechanisms of PTEN loss in cancer: It’s all about diversity. Seminars in Cancer Biology 59, 66–79 (2019).

16. Brennan, C. W. et al. The somatic genomic landscape of glioblastoma. Cell 155, 462–77 (2013).

17. Pulido, R. et al. Precise immunodetection of PTEN protein in human neoplasia. Cold Spring Harbor Perspectives in Medicine 9, 1–30 (2019).

18. Zhang, Y. et al. A Pan-Cancer Proteogenomic Atlas of PI3K/AKT/mTOR Pathway Alterations. Cancer Cell 31, 820-832.e3 (2017).

19. Berger, M. F. et al. The genomic complexity of primary human prostate cancer. Nature 470, 214–220 (2011).

20. Vidotto, T., Tiezzi, D. G. & Squire, J. A. Distinct subtypes of genomic PTEN deletion size influence the landscape of aneuploidy and outcome in prostate cancer. Molecular Cytogenetics 11, 1 (2018).

21. Fan, C., Zhao, C., Shugen Li, F. W. & Wang, J. Significance of PTEN mutation in cellular process, prognosis, and drug selection in clear cell renal cell carcinoma. Frontiers in Oncology 9, 1–9 (2019).

22. Sun, J., Li, S., Wang, F., Fan, C. & Wang, J. Identification of key pathways and genes in pten mutation prostate cancer by bioinformatics analysis. BMC Medical Genetics 20, 1–9 (2019).

23. Imada, E. L. et al. Transcriptional landscape of PTEN loss in primary prostate cancer. BMC Cancer 21, 1–13 (2021).

24. Lin, Z. et al. PTEN loss correlates with T cell exclusion across human cancers. BMC Cancer 21, 1–14 (2021).

25. Jamaspishvili, T. et al. Clinical implications of PTEN loss in prostate cancer. Nature Reviews Urology 15, 222–234 (2018).

26. Hocking, C. et al. Can we accurately report PTEN status in advanced colorectal cancer? BMC Cancer 14, 1–7 (2014).

27. Shao, X. et al. Copy number variation is highly correlated with differential gene expression: A pan-cancer study. BMC Medical Genetics 20, 1–14 (2019).

28. Yoshimoto, M. et al. PTEN genomic deletions that characterize aggressive prostate cancer originate close to segmental duplications. Genes, Chromosomes and Cancer 51, 149–160 (2012).

29. da Costa, A. A. B. A. et al. Low PTEN expression is associated with worse overall survival in head and neck squamous cell carcinoma patients treated with chemotherapy and cetuximab. International Journal of Clinical Oncology 20, 282–289 (2015).

30. Nagata, Y. et al. PTEN activation contributes to tumor inhibition by trastuzumab, and loss of PTEN predicts trastuzumab resistance in patients. Cancer Cell 6, 117–127 (2004).

31. Bucheit, A. D. et al. Complete loss of PTEN protein expression correlates with shorter time to brain metastasis and survival in stage IIIB/C melanoma patients with BRAFV600 mutations. Clinical Cancer Research 20, 5527–5536 (2014).

32. Tachibana, M. et al. Expression and Prognostic Significance of PTEN Product Protein in Patients with Esophageal Squamous Cell Carcinoma. Cancer 94, 1955–1960 (2002).

33. Sawai, H. et al. Loss of PTEN expression is associated with colorectal cancer liver metastasis and poor patient survival. BMC Gastroenterology 8, 1–7 (2008).

34. Sircar, K. et al. PTEN genomic deletion is associated with p-Akt and AR signalling in poorer outcome, hormone refractory prostate cancer. The Journal of pathology 218, 505–513 (2009).

35. Bazzichetto, C. et al. Pten as a prognostic/predictive biomarker in cancer: An unfulfilled promise? Cancers 11, 1–22 (2019).

36. Poluri, R. T. K. & Audet-Walsh, É. Genomic deletion at 10q23 in prostate cancer: More than PTEN loss? Frontiers in Oncology 8, 1–7 (2018).

37. Fusco, N. et al. Pten alterations and their role in cancer management: Are we making headway on precision medicine? Genes 11, 1–19 (2020).

38. Carnero, A. & Paramio, J. M. The PTEN/PI3K/AKT Pathway in vivo, cancer mouse models. Frontiers in Oncology 4, 1–10 (2014).

39. Wang, L. H., Wu, C. F., Rajasekaran, N. & Shin, Y. K. Loss of tumor suppressor gene function in human cancer: An overview. Cellular Physiology and Biochemistry 51, 2647–2693 (2018).

40. Mighell, T. L., Thacker, S., Fombonne, E., Eng, C. & O’Roak, B. J. An Integrated Deep-Mutational-Scanning Approach Provides Clinical Insights on PTEN Genotype-Phenotype Relationships. The American Journal of Human Genetics 106, 818–829 (2020).

41. Hou, S. Q., Ouyang, M., Brandmaier, A., Hao, H. & Shen, W. H. PTEN in the maintenance of genome integrity: From DNA replication to chromosome segregation. BioEssays 39, 1–9 (2017).

42. Salk, J. J., Fox, E. J. & Loeb, L. A. Mutational Heterogeneity in Human Cancers: Origin and Consequences. Annual Review of Pathology: Mechanisms of Disease 5, 51–75 (2010).

43. Lawrence, M. S. et al. Comprehensive genomic characterization of head and neck squamous cell carcinomas. Nature 517, 576–582 (2015).

44. Chen, Z. et al. Crucial role of p53-dependent cellular senescence in suppression of Pten-deficient tumorigenesis. Nature 436, 725–730 (2005).

45. Vahabi, M. et al. MiR-96-5p targets PTEN expression affecting radio-chemosensitivity of HNSCC cells. Journal of Experimental and Clinical Cancer Research 38, 1–16 (2019).

46. Izumi, H. et al. Pathway-specific genome editing of PI3K/mTOR tumor suppressor genes reveals that PTEN loss contributes to cetuximab resistance in head and neck cancer. Molecular Cancer Therapeutics 19, 1562–1571 (2020).

47. Bian, Y. et al. Loss of TGF-β signaling and PTEN promotes head and neck squamous cell carcinoma through cellular senescence evasion and cancer-related inflammation. Oncogene 31, 3322–3332 (2012).

48. dos Santos, L. V., Abrahão, C. M. & William, W. N. Overcoming Resistance to Immune Checkpoint Inhibitors in Head and Neck Squamous Cell Carcinomas. Frontiers in Oncology 11, 1–16 (2021).

49. Serebriiskii, I. G. et al. Comprehensive characterization of PTEN mutational profile in a series of 34,129 colorectal cancers. Nature Communications 13, (2022).

50. Song, M. et al. PTEN Loss Increases PD-L1 Protein Expression and Affects the Correlation between PD-L1 Expression and Clinical Parameters in Colorectal Cancer. PLoS ONE 8, (2013).

51. Ugai, T. et al. Association of PIK3CA mutation and PTEN loss with expression of CD274 (PD-L1) in colorectal carcinoma. OncoImmunology 10, 1956173 (2021).

52. Lastwika, K. J. et al. Control of PD-L1 expression by oncogenic activation of the AKT-mTOR pathway in non-small cell lung cancer. Cancer Research 76, 227–238 (2015).

53. Zhao, J. et al. Immune and genomic correlates of response to anti-PD-1 immunotherapy in glioblastoma. Nature Medicine 25, 462–469 (2019).

54. Imada, E. L. et al. Transcriptional landscape of PTEN loss in primary prostate cancer. bioRxiv 1–42 (2020) doi:10.1101/2020.10.08.332049.

55. Vidotto, T._; et al. PTEN-deficient prostate cancer is associated with an immunosuppressive tumor microenvironment mediated by increased expression of IDO1 and infiltrating FoxP3+ T regulatory cells. The Prostate 1, 1–11 (2019).

56. Mermel, C. H. et al. GISTIC2.0 facilitates sensitive and confident localization of the targets of focal somatic copy-number alteration in human cancers. Genome Biology 12, 1–14 (2011).

57. Zack, T. I. et al. Pan-cancer patterns of somatic copy number alteration. Nature Genetics 45, 1134–1140 (2013).

58. Collado-Torres, L. et al. Reproducible RNA-seq analysis using recount2. Nature Biotechnology 35, 319–321 (2017).

59. Imada, E. L. et al. Recounting the FANTOM CAGE-Associated Transcriptome. Genome Research 30, 1073–1081 (2020).

60. Love, M. I., Huber, W. & Anders, S. Moderated estimation of fold change and dispersion for RNA-seq data with DESeq2. Genome Biology 15, 1–21 (2014).

61. Zhu, A., Ibrahim, J. G. & Love, M. I. Heavy-Tailed prior distributions for sequence count data: Removing the noise and preserving large differences. Bioinformatics 35, 2084–2092 (2019).

62. Liberzon, A. et al. The Molecular Signatures Database Hallmark Gene Set Collection. Cell Systems 1, 417–425 (2015).

63. Wu, T. et al. clusterProfiler 4.0: A universal enrichment tool for interpreting omics data. The Innovation 2, 100141 (2021).

64. Wickham, H. ggplot2: Elegant Graphics for Data Analysis. Springer-Verlag New York. Media vol. 35 (2016).

65. Thorsson, V. et al. The Immune Landscape of Cancer. Immunity 48, 812-830.e14 (2018).

66. Carter, S. L. et al. Absolute quantification of somatic DNA alterations in human cancer. Nature Biotechnology 30, 413–421 (2012).

67. Wellenstein, M. D. & de Visser, K. E. Cancer-Cell-Intrinsic Mechanisms Shaping the Tumor Immune Landscape. Immunity 48, 399–416 (2018).

68. Newman, A. M. et al. Robust enumeration of cell subsets from tissue expression profiles. Nature Methods 1–10 (2015) doi:10.1038/nmeth.3337.

