## Supplementary figures 1 to 15 for "Pan-cancer genomic analysis shows hemizygous *PTEN* loss tumors are associated with immune evasion and poor outcome"

Supplementary Figure 1

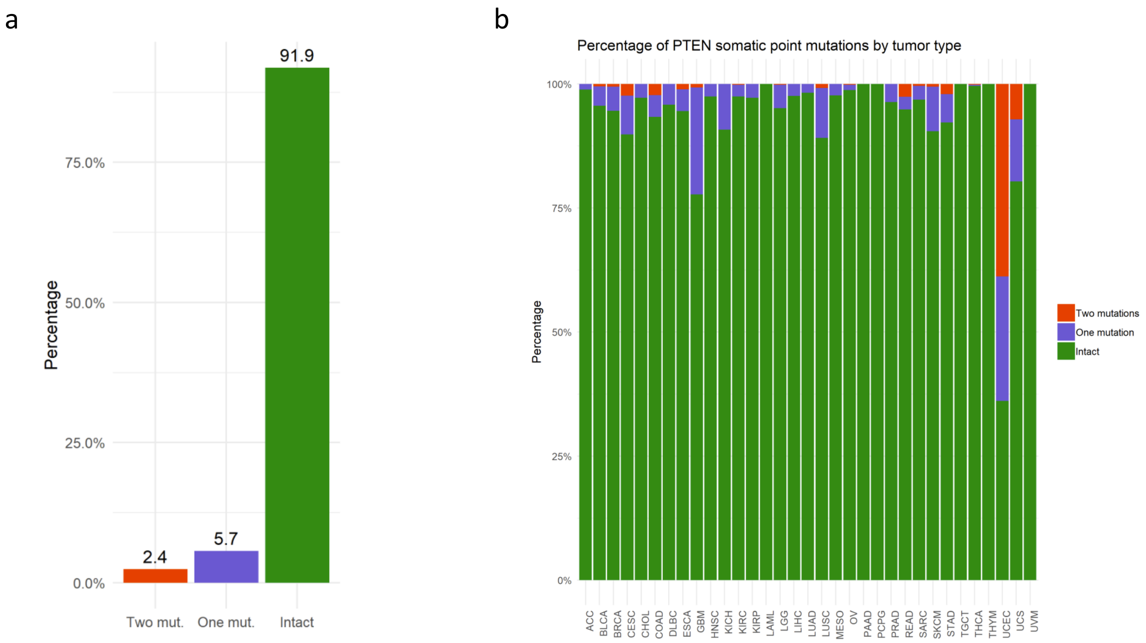


**Supplementary Figure 1.** *PTEN* somatic mutation data from TCGA. (A) PTEN somatic mutation in PanCancer cohort, b) *PTEN* somatic mutations by tumor type. Plotted using *ggplot2* (https://ggplot2.tidyverse.org/).

Supplementary Figure 2

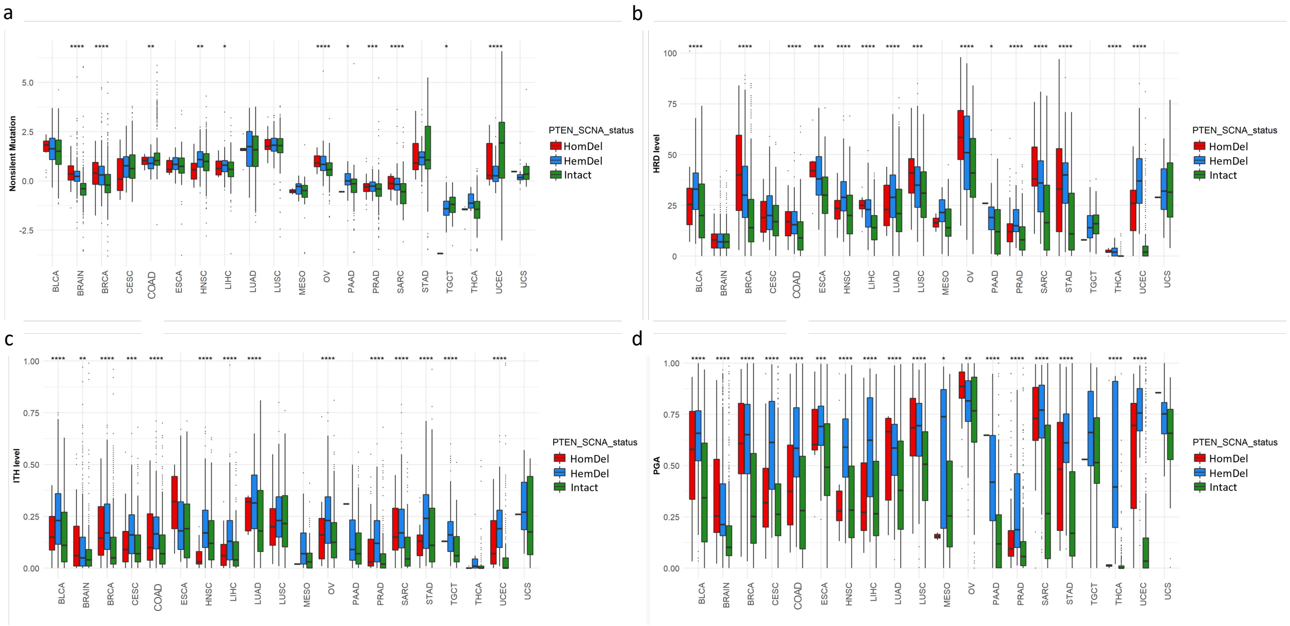


**Supplementary Figure 2.** *PTEN* deletions are associated with genomic alterations in each tumor type. (**a**) Nonsilent mutations. (**b**) Intratumor heterogeneity. (**c**) Homologous repair defects. (**d**) Percent genome altered. Plotted using *ggplot2* (https://ggplot2.tidyverse.org/).

Supplementary Figure 3

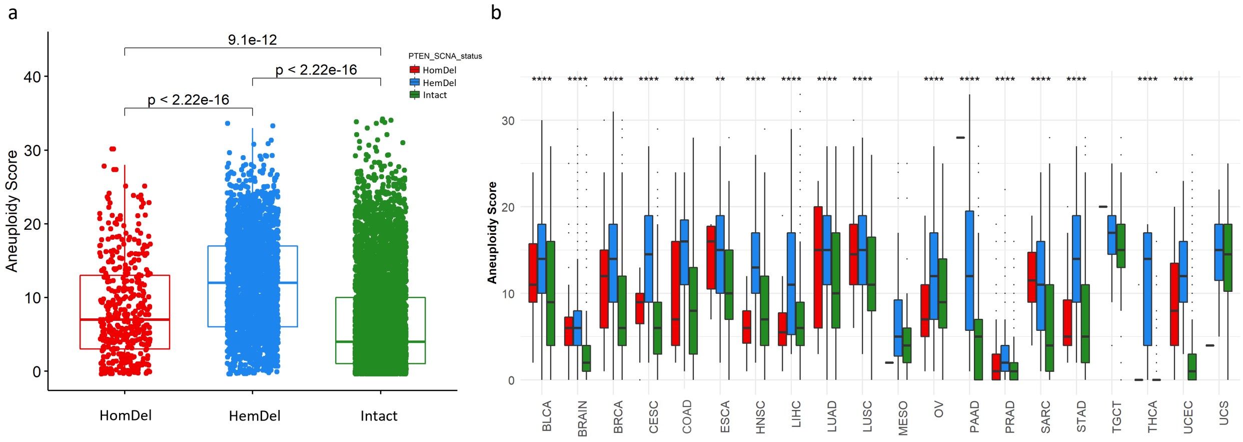


**Supplementary Figure 3.** Association between aneuploidy levels and *PTEN* inactivation status. (**a**) *PTEN* inactivation is significantly associated with increased levels of aneuploidy. (**b**) Aneuploidy score stratified by tumor type. Prostate and brain tumors showed the lowest levels of aneuploidy across tumor types. Aneuploidy levels were determined based on the length of copy number alterations and normalized by tumor type. The scores range from 0, which indicates low levels of chromosome aberrations (copy number gains and losses). On the other hand, high levels of aneuploidy indicate that the genome of these tumors is highly altered, with larger chromosome gains and losses occurring. Plotted using *ggplot2* (https://ggplot2.tidyverse.org/).

Supplementary Figure 4

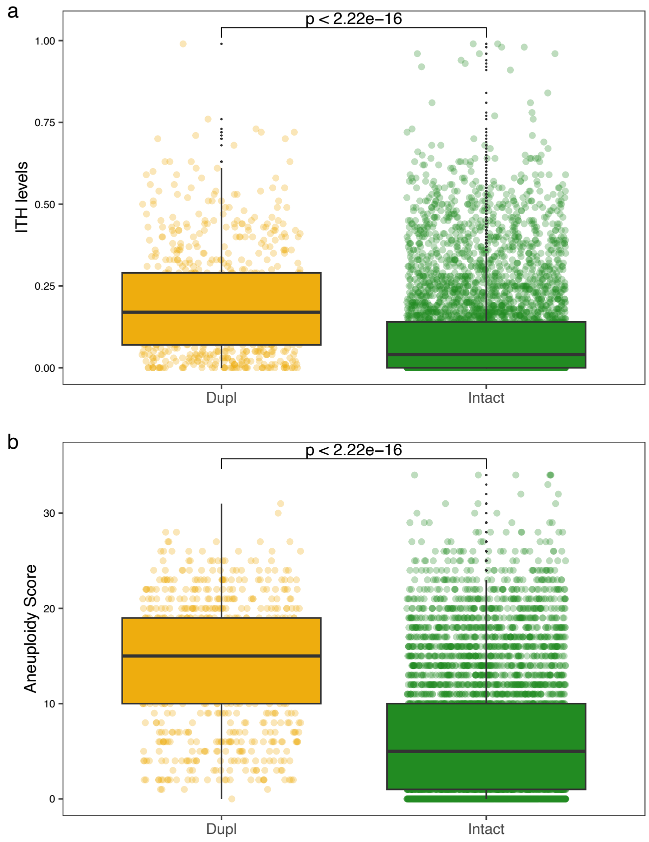


**Supplementary Figure 4.** *PTEN* duplication is associated with genomic alterations. (**a**) *PTEN* Duplication is associated with elevated levels of Intratumor heterogeneity and (**b**) Aneuploidy score. Aneuploidy levels were determined based on the length of copy number alterations and normalized by tumor type. The scores range from 0, which indicates low levels of chromosome aberrations (copy number gains and losses). On the other hand, high levels of aneuploidy indicate that the genome of these tumors is highly altered, with larger chromosome gains and losses occurring. Plotted using *ggplot2* (https://ggplot2.tidyverse.org/).

Supplementary Figure 5

a

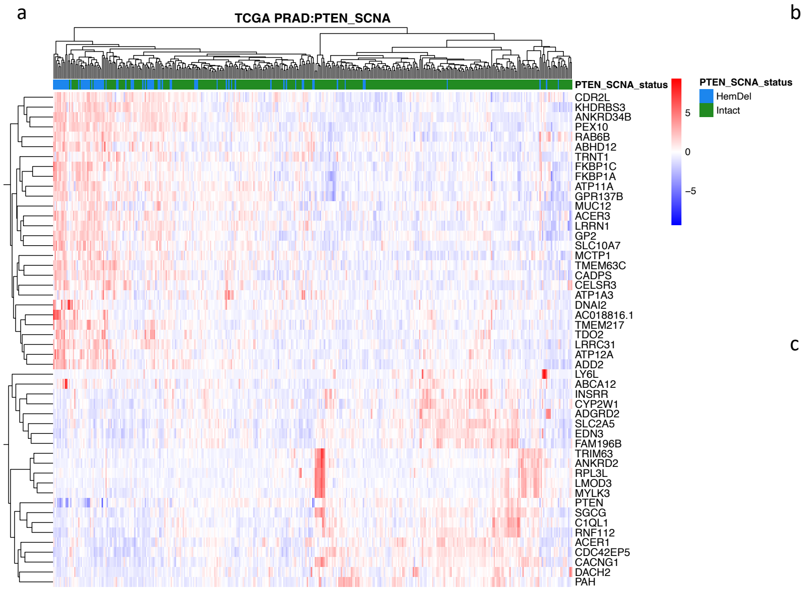


b

| ID | Description | qvalue |
|---|---|---|
| GO:0007018 | microtubule-based movement | 3,41E+11 |
| GO:0007059 | chromosome segregation | 3,41E+11 |
| GO:0000070 | mitotic sister chromatid segregation | 3,41E+11 |
| GO:0098813 | nuclear chromosome segregation | 0.0011 |
| GO:1902850 | microtubule cytoskeleton organization involved in mitosis | 0.0015 |
| GO:0034329 | cell junction assembly | 0.0018 |
| GO:0050808 | synapse organization | 0.0028 |
| GO:0032886 | regulation of microtubule-based process | 0.0038 |
| GO:0000280 | nuclear division | 0.0065 |
| GO:0007156 | homophilic cell adhesion via plasma membrane adhesion molecules | 0.0072 |

c

| ID | Description | qvalue |
|---|---|---|
| GO:0002181 | cytoplasmic translation | 3,03E+07 |
| GO:0006936 | muscle contraction | 1,28E+10 |
| GO:0016054 | organic acid catabolic process | 1,86E+11 |
| GO:0044282 | small molecule catabolic process | 8,90E+11 |
| GO:0008015 | blood circulation | 0.0020 |
| GO:0086005 | ventricular cardiac muscle cell action potential | 0.0070 |
| GO:0010273 | detoxification of copper ion | 0.0154 |
| GO:1990169 | stress response to copper ion | 0.0154 |
| GO:0061687 | detoxification of inorganic compound | 0.0174 |
| GO:0086009 | membrane repolarization | 0.0174 |

**Supplementary Figure 5.** *PTEN* HemDel PRAD showed distinct transcriptome activity. (**a**) Transcriptome heatmap exhibiting clustering of top 50 DEGs (Log2Fold change >0.58). (**b**) Enriched GO (BP) pathways of the upregulated genes. (**c**) Enriched GO (BP) pathways of the downregulated genes. The color scale in the heatmap represents the Z-score of the normalized read counts for each gene, where the red scale color indicates upregulated and blue low-expressed genes. Enrichment analyses were performed using *clusterProfiler* and *p*-adjusted value = 0.05 as the cutoff. Tables show the top 10 most significant Gene Ontology (BP) terms. Plotted using *pheatmap* (https://rdocumentation.org/packages/pheatmap/versions/1.0.12).

Supplementary Figure 6

a

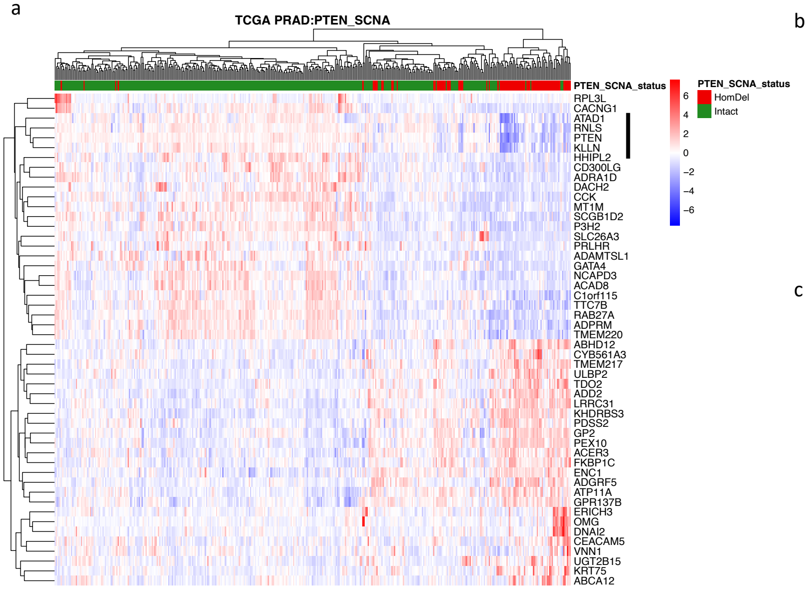


b

| ID | Description | qvalue |
|---|---|---|
| GO:0044770 | cell cycle phase transition | 1,09E+12 |
| GO:0007059 | chromosome segregation | 1,39E+12 |
| GO:0007018 | microtubule-based movement | 2,92E+11 |
| GO:1901987 | regulation of cell cycle phase transition | 4,63E+12 |
| GO:0006281 | DNA repair | 6,03E+11 |
| GO:0000819 | sister chromatid segregation | 0.0017 |
| GO:0000070 | mitotic sister chromatid segregation | 0.0017 |
| GO:0044782 | cilium organization | 0.0017 |
| GO:0001578 | microtubule bundle formation | 0.0020 |
| GO:0048285 | organelle fission | 0.0033 |

c

| ID | Description | qvalue |
|---|---|---|
| GO:0008015 | blood circulation | 8,75E+09 |
| GO:0006936 | muscle contraction | 1,16E+11 |
| GO:0003018 | vascular process in circulatory system | 3,47E+11 |
| GO:0034765 | regulation of ion transmembrane transport | 4,05E+11 |
| GO:0099177 | regulation of trans-synaptic signaling | 0.0011 |
| GO:0015844 | monoamine transport | 0.0022 |
| GO:0008016 | regulation of heart contraction | 0.0038 |
| GO:0016054 | organic acid catabolic process | 0.0041 |
| GO:0072503 | cellular divalent inorganic cation homeostasis | 0.0041 |
| GO:0007188 | adenylate cyclase-modulating G protein-coupled receptor signaling pathway | 0.0041 |

**Supplementary Figure 6.** *PTEN* HomDel PRAD showed distinct transcriptome activity. **a)** Transcriptome heatmap exhibiting clustering of top 50 DEGs (Log2Fold change >0.58). **b)** Enriched GO (BP) pathways of the upregulated genes. **c)** Enriched GO (BP) pathways of the downregulated genes. The color scale in the heatmap represents the Z-score of the normalized read counts for each gene, where the red scale color indicates upregulated and blue low-expressed genes. Enrichment analyses were performed using *clusterProfiler* and *p*-adjusted value = 0.05 as the cutoff. Tables show the top 10 most significant Gene Ontology (BP) terms. Vertical bar indicates the closely linked passenger genes (*ATAD1, RNLS* and *KLLN*) that are concurrently down-regulated when *PTEN* is lost by a large deletion of chromosome 10. Plotted using *pheatmap* (https://rdocumentation.org/packages/pheatmap/versions/1.0.12).

Supplementary Figure 7

a

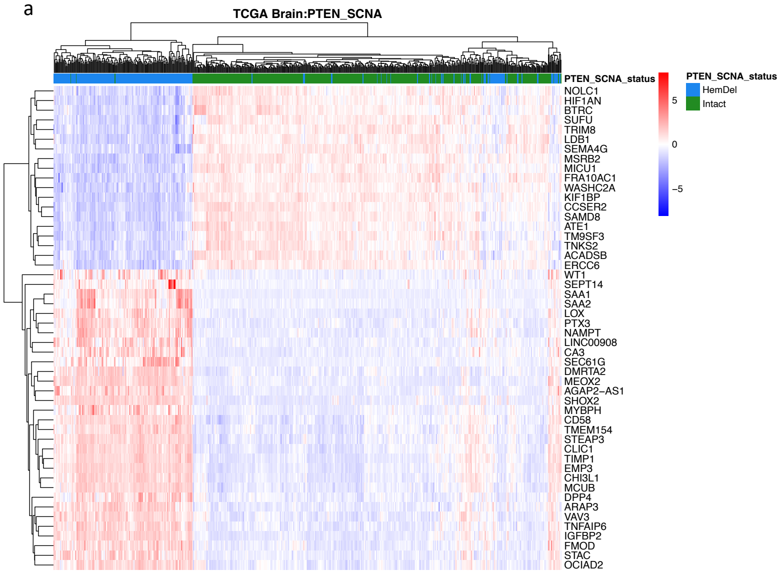


b

| ID | Description | qvalue |
|---|---|---|
| GO:0002250 | adaptive immune response | 9,17E-10 |
| GO:0042110 | T cell activation | 3,96E-06 |
| GO:0030198 | extracellular matrix organization | 5,64E-05 |
| GO:0002443 | leukocyte mediated immunity | 2,69E-01 |
| GO:0002237 | response to molecule of bacterial origin | 4,85E-01 |
| GO:0045785 | positive regulation of cell adhesion | 6,54E-02 |
| GO:1903037 | regulation of leukocyte cell-cell adhesion | 7,12E-02 |
| GO:0007159 | leukocyte cell-cell adhesion | 1,31E+00 |
| GO:0050900 | leukocyte migration | 1,39E-01 |
| GO:0050863 | regulation of T cell activation | 2,64E+00 |

c

| ID | Description | qvalue |
|---|---|---|
| GO:0022613 | ribonucleoprotein complex biogenesis | 1,67E+08 |
| GO:0008380 | RNA splicing | 1,67E+08 |
| GO:0050804 | modulation of chemical synaptic transmission | 2,70E+10 |
| GO:0099003 | vesicle-mediated transport in synapse | 6,22E+09 |
| GO:0022618 | ribonucleoprotein complex assembly | 8,34E+09 |
| GO:0071826 | ribonucleoprotein complex subunit organization | 1,97E+11 |
| GO:0050808 | synapse organization | 5,09E+10 |
| GO:0016197 | endosomal transport | 5,10E+11 |
| GO:0043543 | protein acylation | 7,89E+10 |
| GO:0001505 | regulation of neurotransmitter levels | 8,50E+11 |

**Supplementary Figure 7.** *PTEN* HemDel Brain (GBM + LGG) showed distinct transcriptome activity. (**a**) Transcriptome heatmap exhibiting clustering of top 50 DEGs (Log2Fold change >0.58). (**b**) Enriched GO (BP) pathways of the upregulated genes. (**c**) Enriched GO (BP) pathways of the downregulated genes. The color scale in the heatmap represents the Z-score of the normalized read counts for each gene, where the red scale color indicates upregulated and blue low-expressed genes. Enrichment analyses were performed using *clusterProfiler* and *p*-adjusted value = 0.05 as the cutoff. Tables show the top 10 most significant Gene Ontology (BP) terms. Plotted using *pheatmap* (https://rdocumentation.org/packages/pheatmap/versions/1.0.12).

Supplementary Figure 8

a

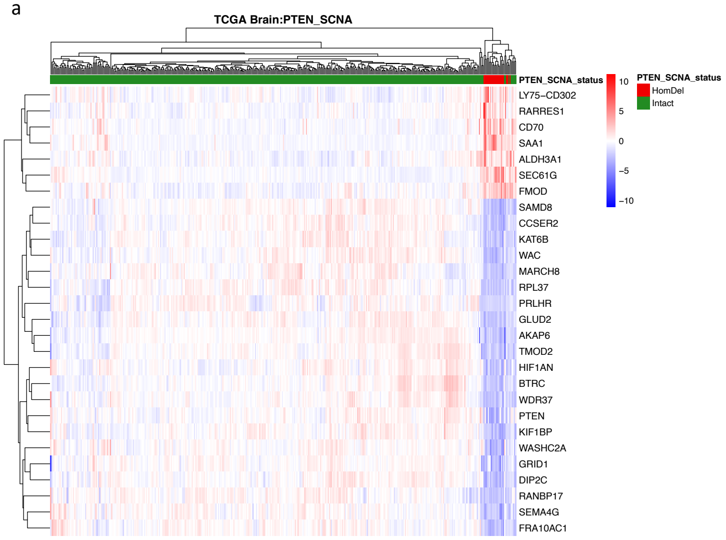


b

| ID | Description | qvalue |
|---|---|---|
| GO:0002250 | adaptive immune response | 4,71E-13 |
| GO:0042110 | T cell activation | 2,76E-09 |
| GO:0050867 | positive regulation of cell activation | 3,49E-08 |
| GO:0002443 | leukocyte mediated immunity | 7,82E-08 |
| GO:1903037 | regulation of leukocyte cell-cell adhesion | 2,31E-06 |
| GO:0050778 | positive regulation of immune response | 3,60E-05 |
| GO:0007159 | leukocyte cell-cell adhesion | 7,56E-06 |
| GO:0019221 | cytokine-mediated signaling pathway | 7,58E-06 |
| GO:0050870 | positive regulation of T cell activation | 1,71E-02 |
| GO:0001819 | positive regulation of cytokine production | 1,10E-01 |

c

| ID | Description | qvalue |
|---|---|---|
| GO:0050804 | modulation of chemical synaptic transmission | 1,60E+01 |
| GO:0099177 | regulation of trans-synaptic signaling | 1,60E+01 |
| GO:0099504 | synaptic vesicle cycle | 2,43E+03 |
| GO:0050808 | synapse organization | 9,90E+04 |
| GO:0006397 | mRNA processing | 2,35E+08 |
| GO:0016079 | synaptic vesicle exocytosis | 2,35E+08 |
| GO:0060078 | regulation of postsynaptic membrane potential | 7,11E+07 |
| GO:0099643 | signal release from synapse | 1,55E+09 |
| GO:0016358 | dendrite development | 5,48E+08 |
| GO:0001505 | regulation of neurotransmitter levels | 1,31E+09 |

**Supplementary Figure 8.** *PTEN* HomDel Brain (GBM + LGG) showed distinct transcriptome activity. (**a**) Transcriptome heatmap exhibiting clustering of top 50 DEGs (Log2Fold change >0.58). Of the 4106 overexpressed genes in the HomDel tumors 66.7% were also overexpressed in the 3562 HemDel tumors. Similarly, of the 7332 underexpressed genes in the HomDel tumors 59.9% were also underexpressed in the 10902 underexpressed genes in the HemDel tumors (**b**) Enriched GO (BP) pathways of the upregulated genes. The color scale in the heatmap represents the Z-score of the normalized read counts for each gene, where the red scale color indicates upregulated and blue low-expressed genes. Enrichment analyses were performed using *clusterProfiler* and *p*-adjusted value = 0.05 as the cutoff. Tables show the top 10 most significant Gene Ontology (BP) terms. Plotted using *pheatmap* (https://rdocumentation.org/packages/pheatmap/versions/1.0.12).

Supplementary Figure 9

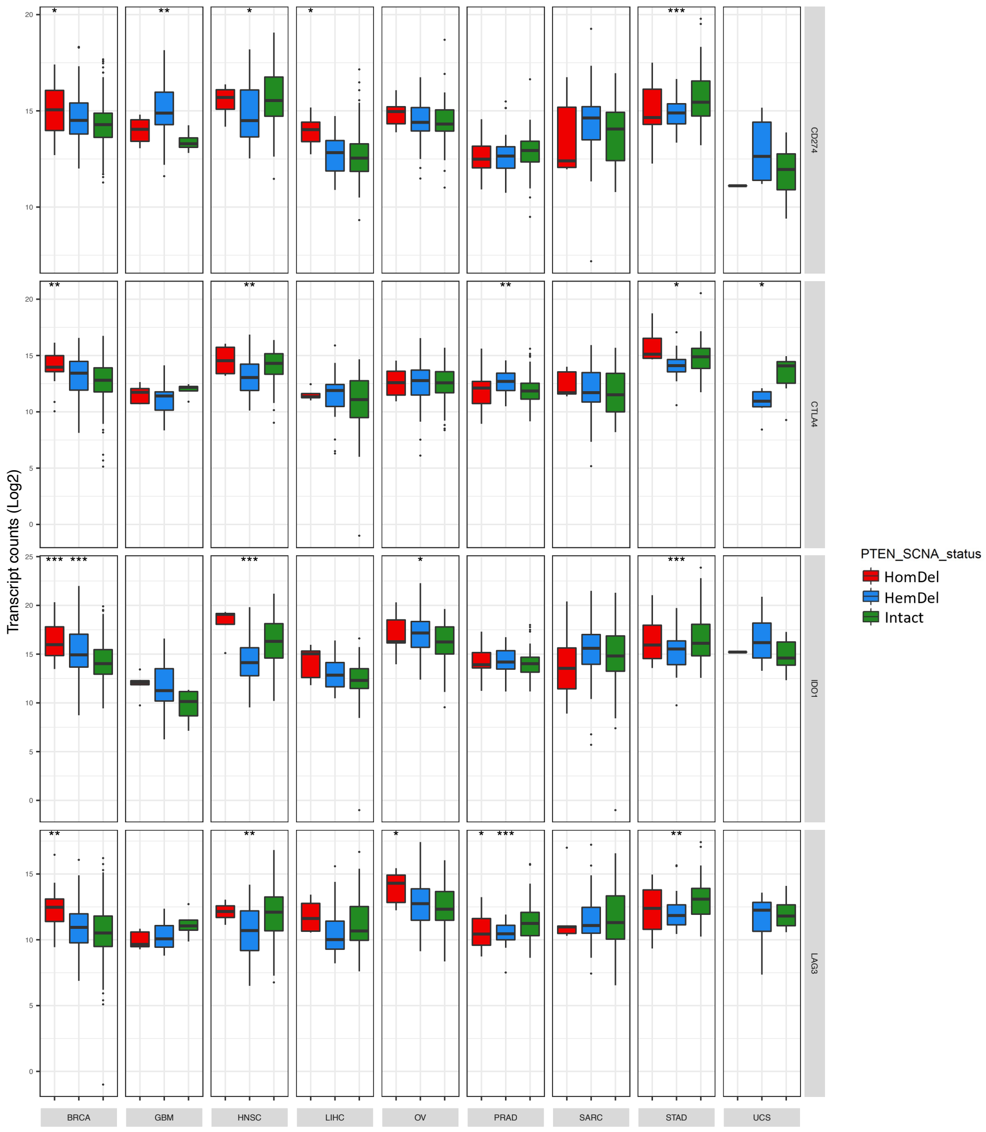


**Supplementary Figure 9.** The RNA-sequencing expression level of immunomodulatory genes *CD274* (*PD-L1*), *CTLA4*, *IDO1*, and *LAG3* in BRCA, GBM, HNSC, LIHC, OV, PRAD, SARC, STAD, and UCS. *: *p*-value < 0.05; **: *p*-value < 0.01, by Mann-Whitney test. *CD274* (*PD-L1*), Programmed Cell Death 1 Ligand 1; *CTLA4*, Cytotoxic T-Lymphocyte Associated Protein 4; *IDO1*, Indoleamine 2,3-Dioxygenase 1; LAG3, Lymphocyte Activating 3. Plotted using *ggplot2* (https://ggplot2.tidyverse.org/).

Supplementary Figure 10

a

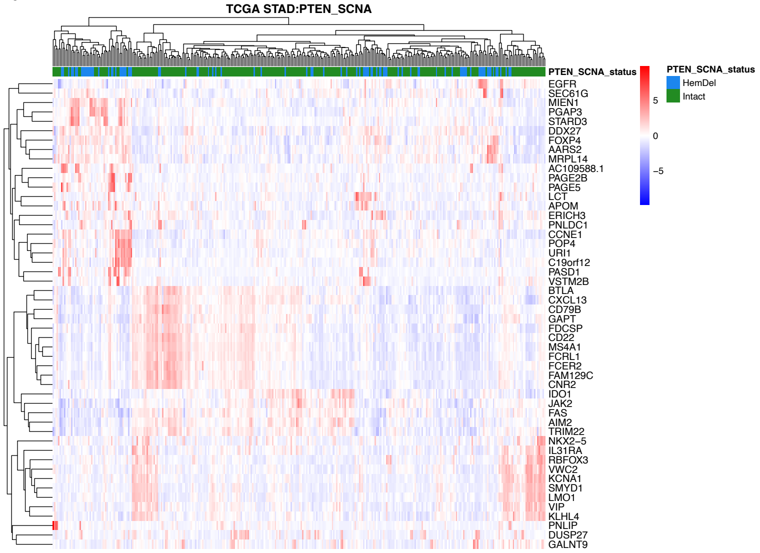


b

| ID | Description | qvalue |
|---|---|---|
| GO:0000819 | sister chromatid segregation | 1,75E+06 |
| GO:0000070 | mitotic sister chromatid segregation | 1,75E+06 |
| GO:0007059 | chromosome segregation | 2,77E+07 |
| GO:0000280 | nuclear division | 2,77E+07 |
| GO:0098813 | nuclear chromosome segregation | 5,81E+06 |
| GO:0051321 | meiotic cell cycle | 1,05E+12 |
| GO:0016126 | sterol biosynthetic process | 5,01E+11 |
| GO:0006695 | cholesterol biosynthetic process | 5,01E+11 |
| GO:1902653 | secondary alcohol biosynthetic process | 5,01E+11 |
| GO:0034660 | ncRNA metabolic process | 7,25E+11 |

c

| ID | Description | qvalue |
|---|---|---|
| GO:0002250 | adaptive immune response | 1,55E-22 |
| GO:0051249 | regulation of lymphocyte activation | 5,34E-12 |
| GO:0042110 | T cell activation | 7,73E-12 |
| GO:0050778 | positive regulation of immune response | 9,10E-11 |
| GO:0002443 | leukocyte mediated immunity | 8,74E-09 |
| GO:0007159 | leukocyte cell-cell adhesion | 3,17E-07 |
| GO:1903037 | regulation of leukocyte cell-cell adhesion | 4,46E-07 |
| GO:0050867 | positive regulation of cell activation | 7,92E-07 |
| GO:0070661 | leukocyte proliferation | 4,32E-06 |
| GO:0046651 | lymphocyte proliferation | 1,44E-04 |

**Supplementary Figure 10.** *PTEN* HemDel STAD showed distinct transcriptome activity. (**a**) Transcriptome heatmap exhibiting clustering of top 50 DEGs (Log2Fold change >0.58). (**b**) Enriched GO (BP) pathways of the upregulated genes. (**c**) Enriched GO (BP) pathways of the downregulated genes. The color scale in the heatmap represents the Z-score of the normalized read counts for each gene, where the red scale color indicates upregulated and blue low-expressed genes. Enrichment analyses were performed using *clusterProfiler* and *p*-adjusted value = 0.05 as the cutoff. Tables show the top 10 most significant Gene Ontology (BP) terms. Plotted using *pheatmap* (https://rdocumentation.org/packages/pheatmap/versions/1.0.12).

Supplementary Figure 11

a

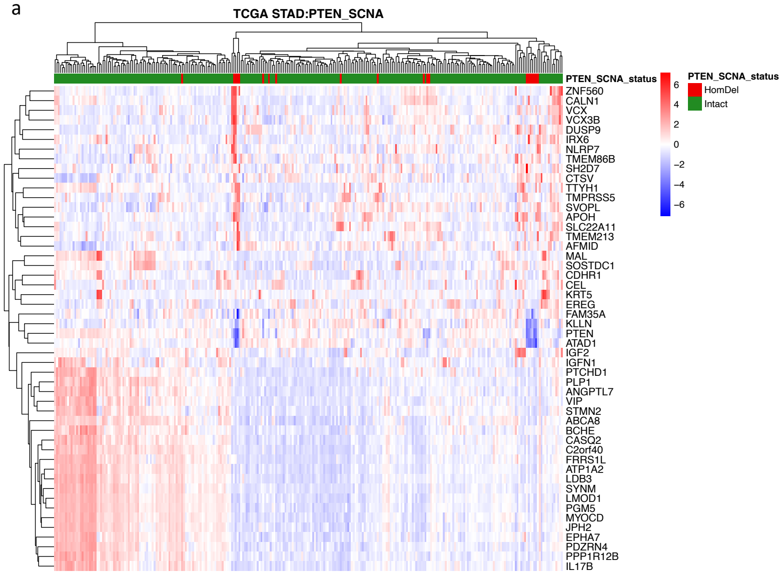


b

| ID | Description | qvalue |
|---|---|---|
| GO:0003012 | muscle system process | 5,90E-07 |
| GO:0051480 | regulation of cytosolic calcium ion concentration | 7,86E+02 |
| GO:0008015 | blood circulation | 2,35E+04 |
| GO:0010959 | regulation of metal ion transport | 7,72E+05 |
| GO:0072507 | divalent inorganic cation homeostasis | 7,96E+05 |
| GO:0008016 | regulation of heart contraction | 2,15E+08 |
| GO:0042692 | muscle cell differentiation | 3,11E+07 |
| GO:0060402 | calcium ion transport into cytosol | 5,62E+08 |
| GO:0034765 | regulation of ion transmembrane transport | 7,90E+07 |
| GO:0006816 | calcium ion transport | 1,21E+09 |

**Supplementary Figure 11.** *PTEN* HomDel STAD showed distinct transcriptome activity. (**a**) Transcriptome heatmap exhibiting clustering of top 50 DEGs (Log2Fold change >0.58). (**b**) Enriched GO (BP) pathways of the upregulated genes. The color scale in the heatmap represents the Z-score of the normalized read counts for each gene, where the red scale color indicates upregulated and blue low-expressed genes. Enrichment analyses were performed using *clusterProfiler* and *p*-adjusted value = 0.05 as the cutoff. Tables show the top 10 most significant Gene Ontology (BP) terms. Plotted using *pheatmap* (https://rdocumentation.org/packages/pheatmap/versions/1.0.12).

Supplementary Figure 12

a

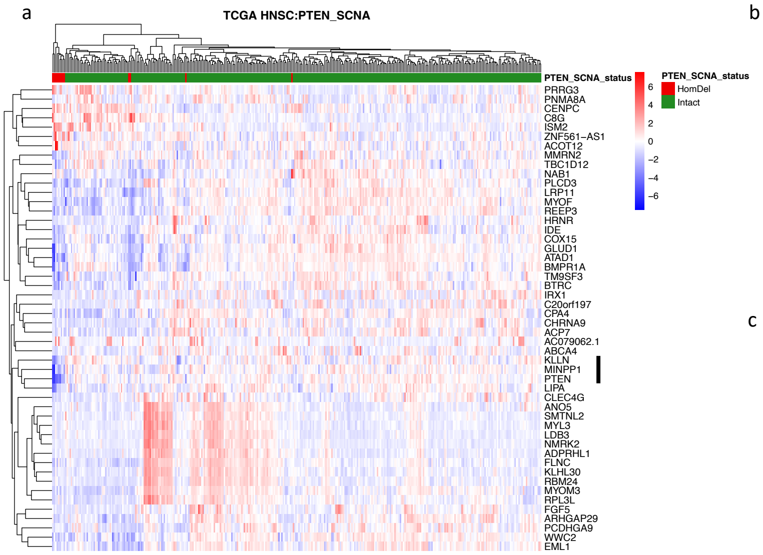


b

| ID | Description | qvalue |
|---|---|---|
| GO:0006260 | DNA replication | 8,25E+09 |
| GO:0007059 | chromosome segregation | 8,25E+09 |
| GO:0051321 | meiotic cell cycle | 5,49E+09 |
| GO:0098813 | nuclear chromosome segregation | 1,39E+12 |
| GO:0000280 | nuclear division | 1,39E+12 |
| GO:0034508 | centromere complex assembly | 1,92E+11 |
| GO:0006335 | DNA replication-dependent nucleosome assembly | 0.0023 |
| GO:0034723 | DNA replication-dependent nucleosome organization | 0.0023 |
| GO:0071824 | protein-DNA complex subunit organization | 0.0028 |
| GO:0007098 | centrosome cycle | 0.0028 |

c

| ID | Description | qvalue |
|---|---|---|
| GO:0003012 | muscle system process | 2,71E-09 |
| GO:0055001 | muscle cell development | 2,11E+00 |
| GO:0014706 | striated muscle tissue development | 3,43E+01 |
| GO:0030239 | myofibril assembly | 7,66E+00 |
| GO:0007517 | muscle organ development | 2,85E+04 |
| GO:0031032 | actomyosin structure organization | 3,33E+04 |
| GO:0048738 | cardiac muscle tissue development | 3,80E+04 |
| GO:0050879 | multicellular organismal movement | 6,46E+04 |
| GO:0050881 | musculoskeletal movement | 6,46E+04 |
| GO:0008015 | blood circulation | 3,51E+06 |

**Supplementary Figure 12.** *PTEN* HomDel HNSC showed distinct transcriptome activity. (**a**) Transcriptome heatmap exhibiting clustering of top 50 DEGs (Log2Fold change >0.58). (**b**) Enriched GO (BP) pathways of the upregulated genes. (**c**) Enriched GO (BP) pathways of the downregulated genes. The color scale in the heatmap represents the Z-score of the normalized read counts for each gene, where the red scale color indicates upregulated and blue low-expressed genes. Enrichment analyses were performed using *clusterProfiler* and *p*-adjusted value = 0.05 as the cutoff. Tables show the top 10 most significant Gene Ontology (BP) terms. Vertical bar indicates the two closely linked passenger genes (*MINPP1 and KLLN*) that are concurrently down-regulated when PTEN is lost by a large deletion of chromosome 10. Plotted using *pheatmap* (https://rdocumentation.org/packages/pheatmap/versions/1.0.12).

Supplementary Figure 13

a

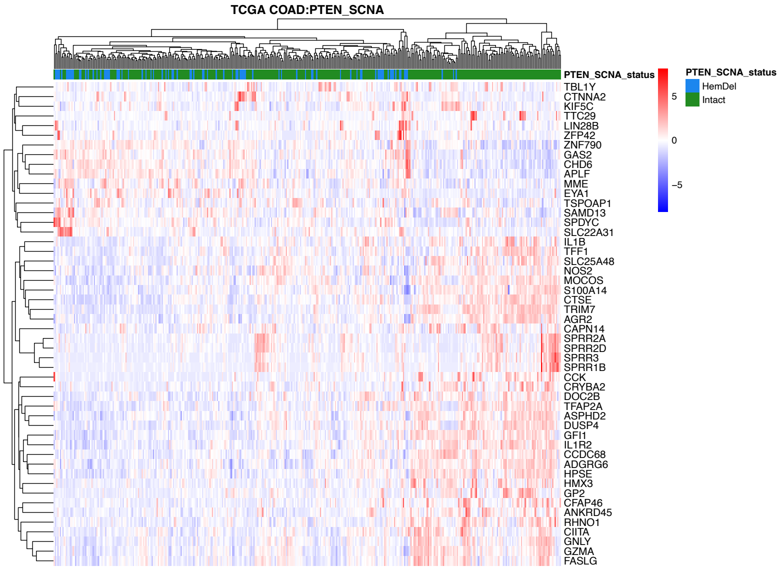


b

| ID | Description | qvalue |
|---|---|---|
| GO:0034341 | response to interferon-gamma | 1,93E+10 |
| GO:0002449 | lymphocyte mediated immunity | 1,93E+10 |
| GO:0002250 | adaptive immune response | 1,93E+10 |
| GO:0002228 | natural killer cell mediated immunity | 8,21E+10 |
| GO:0045333 | cellular respiration | 1,12E+12 |
| GO:0002396 | MHC protein complex assembly | 2,46E+12 |
| GO:0001906 | cell killing | 2,49E+12 |
| GO:0009615 | response to virus | 2,49E+12 |
| GO:0001909 | leukocyte mediated cytotoxicity | 3,67E+12 |
| GO:0002495 | antigen processing and presentation of peptide antigen via MHC class II | 3,67E+12 |

**Supplementary Figure 13.** *PTEN* HemDel COAD showed distinct transcriptome activity. (**a**) Transcriptome heatmap exhibiting clustering of top 50 DEGs (Log2Fold change >0.58). (**b**) Enriched GO (BP) pathways of the downregulated genes. The color scale in the heatmap represents the Z-score of the normalized read counts for each gene, where the red scale color indicates upregulated and blue low-expressed genes. Enrichment analyses were performed using *clusterProfiler* and *p*-adjusted value = 0.05 as the cutoff. Tables show the top 10 most significant Gene Ontology (BP) terms. Plotted using *pheatmap* (https://rdocumentation.org/packages/pheatmap/versions/1.0.12).

Supplementary Figure 14

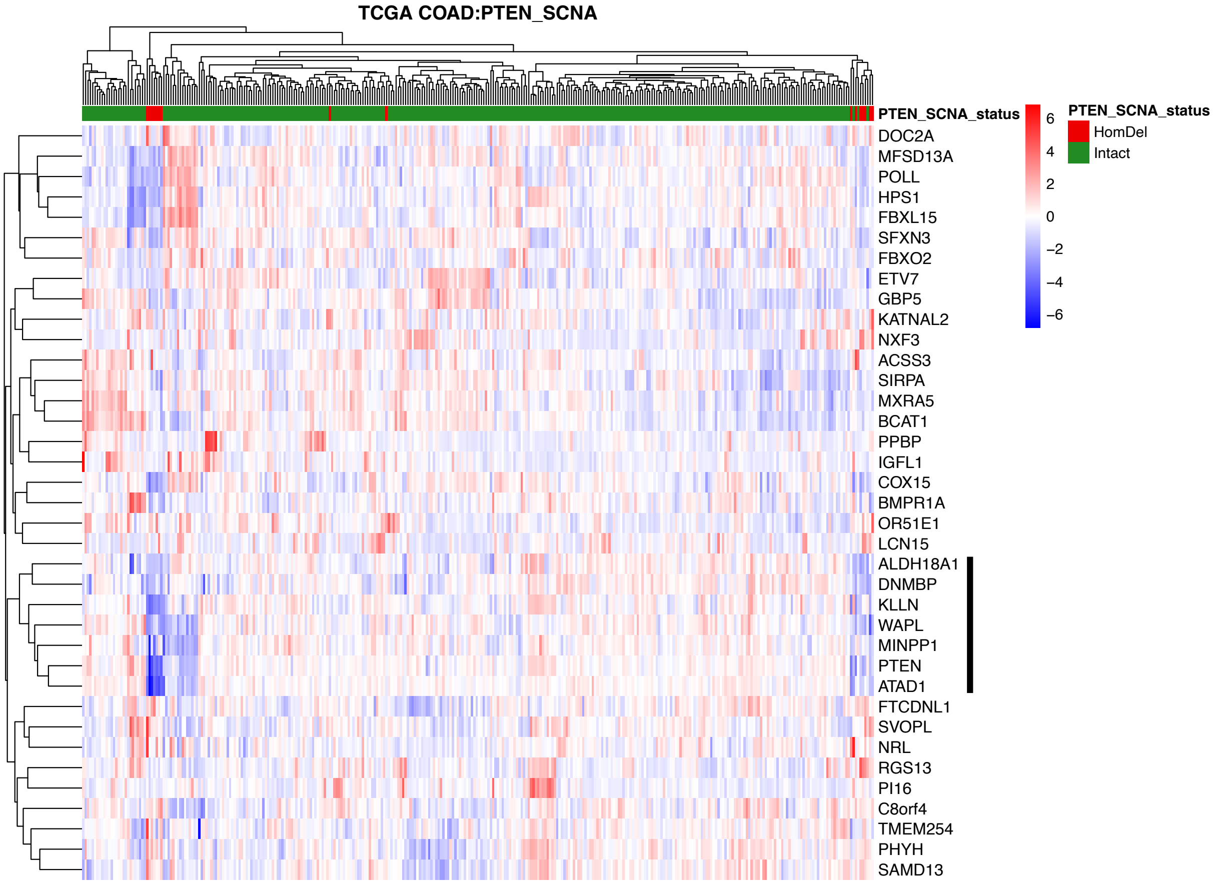


**Supplementary Figure 14.** *PTEN* HomDel COAD showed distinct transcriptome activity. Transcriptome heatmap exhibiting clustering of top 50 DEGs (Log2Fold change >0.58). Vertical bar indicates the closely linked passenger genes (*ALDH18A1, DNMBP, KLLN*, *WAPL, MINPP1, ATAD1)* that are concurrently down-regulated when PTEN is lost by a large deletion of chromosome 10. Plotted using *pheatmap* (https://rdocumentation.org/packages/pheatmap/versions/1.0.12).

Supplementary Figure 15

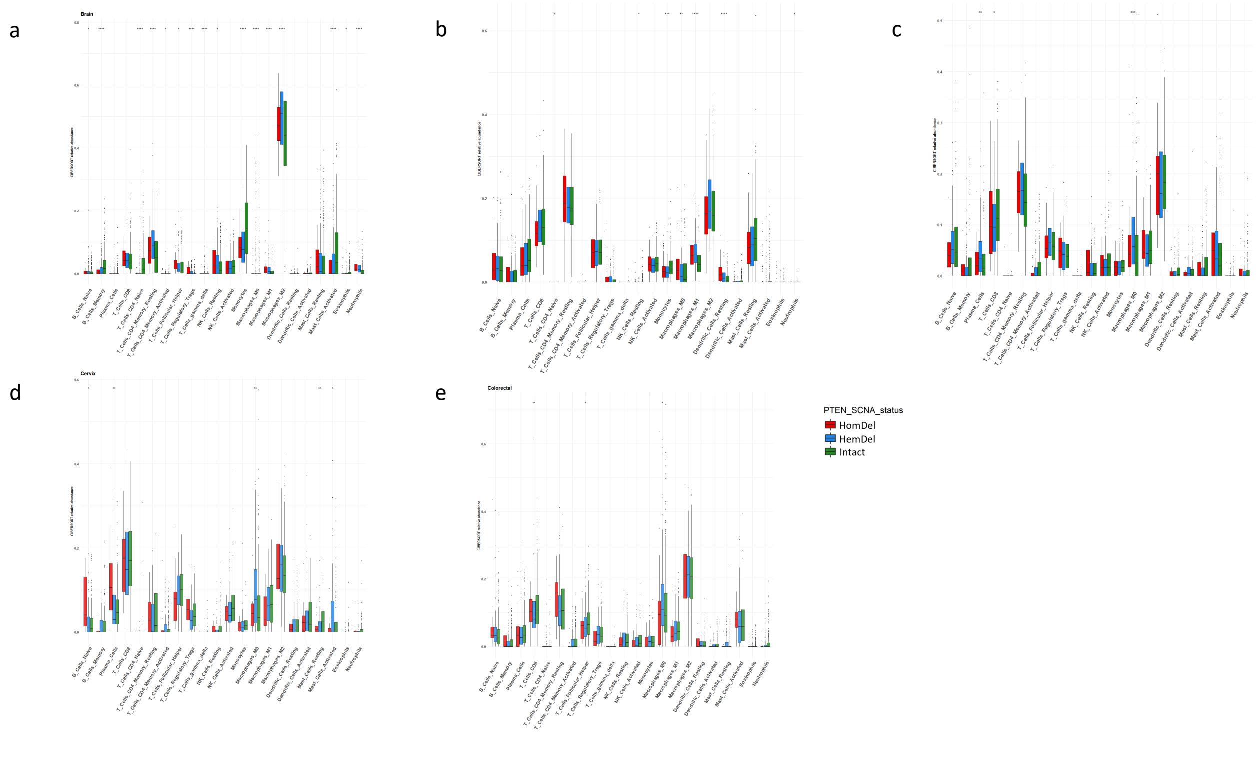


**Supplementary Figure 15.** CIBERSORT-derived immune-cell abundance of 22 distinct cell subsets based on *PTEN* status and tumor type. Y-axis shows CIBERSORT relative scores. (**a**) *PTEN* HemDel in brain tumors was linked to higher CD4 T memory cells and M2 macrophage abundance and lower M1 macrophages, monocytes, and CD8 T cells. (**b**) Prostate tumors with *PTEN* HemDel showed increased M1 macrophages and decreased M0 macrophages and monocytes. (**c**) Stomach tumors with *PTEN* HemDel showed increased plasma and M0 macrophages and decreased in CD8 T cells. (**d**) Cervical tumors with *PTEN* HemDel were linked to higher M0 macrophages and mast cells activated. (**e**) Colorectal tumors with *PTEN* HemDel were linked to higher M0 macrophages and lower CD8 T cells. *: $p$-value < 0.05; **: $p$-value < 0.01, by Mann-Whitney test. DC, dendritic cell; NK, natural-killer cell; PCa, prostate cancer; Treg, regulatory T cell; M0, M0 macrophage; M1, M1 macrophage; M2, M2 macrophage. Plotted using *ggplot2* (https://ggplot2.tidyverse.org/).
