## Supplementary tables 1 to 3 for "Pan-cancer genomic analysis shows hemizygous *PTEN* loss tumors are associated with immune evasion and poor outcome"

Supplementary Table 1

| PTEN_SCNA_status | TP53 Mut | TP53 WT | χ2 | p-value |
|---|---|---|---|---|
| HemDel | 1343 | 1155 | 448.49 | < 0.001 |
| HomDel | 194 | 223 | | |
| Intact | 1786 | 4206 | | |
| **Total** | **3323** | **5584** | | |

**Supplementary Table 1**. A chi-square test of independence showed that there was a significant association between *PTEN* status and somatic mutation status of *TP53*, $\chi 2$ (2, N = 7866) = 448.4, p = < 0.001. *TP53* mutation status was more likely than *TP53* WT in PTEN HemDel.

Supplementary Table 2

a)

| Variable | PanCancer (OS) | |
|---|---|---|
| | HR (95% CI) | p* |
| **PTEN CNA + TP53** | | |
| Intact + WT | - | - |
| Intact + MonAll | 2.07 (1.86 to 2.3) | p<0.001 |
| Intact + BiAll | 1.39 (1.06 to 0.0) | 0.019 |
| HomDel + WT | 2.09 (1.68 to 2.6) | p<0.001 |
| HomDel + MonAll | 1.55 (1.19 to 2.02) | 0.001 |
| HomDel + BiAll | 3.79 (1.7 to 8.46) | 0.001 |
| HemDel + WT | 2.67 (2.41 to 2.95) | p<0.001 |
| HemDel + MonAll | 2.36 (2.12 to 2.63) | p<0.001 |
| HemDel + BiAll | 2.84 (2.17 to 3.72) | p<0.001 |

b)

| Variable | HNSC (OS) | |
|---|---|---|
| | HR (95% CI) | p* |
| **PTEN CNA + TP53** | | |
| Intact + WT | - | - |
| Intact + MonAll | 1.90 (1.29 to 2.81) | 0.001 |
| Intact + BiAll | 1.59 (0.91 to 2.55) | 0.10 |
| HomDel + WT | 1.6 (0.00 to -) | - |
| HomDel + MonAll | 0.78 (0.10 to 5.7) | 0.8 |
| HomDel + BiAll | 4.14 (0.57 to 30.2) | 0.16 |
| HemDel + WT | 1.99 (0.99 to 4.01) | 0.053 |
| HemDel + MonAll | 1.67 (1.08 to 2.6) | 0.023 |
| HemDel + BiAll | 1.47 (0.65 to 3.29) | 0.35 |

**Supplementary Table 2**. Cox Model analysis of overall survivor for the co-occurrence of *TP53* mutation and *PTEN* SCNA. **(A)** *PTEN* HemDel and *TP53* monoallelic mutation tumors are associated with similar hazard ratio levels across the entire TCGA cohort **(B)** Head and neck tumors also showed that *PTEN* HemDel and *TP53* monoallelic mutation exhibited similar hazard ratios. HR, Hazard Ratio; *TP53* MonAll, monoallelic loss for *TP53;* BiAll, biallelic loss for *TP53*.

Supplementary table 3

| Tumor type | HemDel | HomDel | Int | DE Genes (HemDel vs Intact) | | DE Genes (HomDel vs Intact) | |
|---|---|---|---|---|---|---|---|
| | | | | Up | Down | Up | Down |
| Brain | 235 | 22 | 406 | 3562 | 10902 | 4106 | 7332 |
| Colorectal | 86 | 16 | 319 | 1086 | 1536 | 23 | 175 |
| Cervix | 74 | 14 | 186 | 728 | 649 | 743 | 512 |
| Head and Neck | 117 | 13 | 329 | 1727 | 1842 | 444 | 1148 |
| Prostate | 73 | 85 | 324 | 2076 | 2966 | 3304 | 3835 |
| Sarcoma | 120 | 15 | 106 | 2032 | 3725 | 241 | 708 |
| Stomach | 90 | 20 | 256 | 3216 | 2814 | 683 | 1434 |
| | 795 | 185 | 1926 | 14427 | 24432 | 9544 | 15144 |

**Supplementary Table 3**. The number of differentially expressed genes (DEGs) comparing HemDel and HomDel to intact in seven solid tumors. FDR < 0.05. *PTEN* deletion status: HemDel: *PTEN* hemizygous deletion. HomDel: *PTEN* homozygous deletion. Int: *PTEN* intact. FDR, False discovery rate.
